# Preventing multiple resistance above all: new insights for managing fungal adaptation

**DOI:** 10.1101/2022.12.17.520869

**Authors:** Agathe Ballu, Claire Ugazio, Clémentine Duplaix, Alicia Noly, Juerg Wullschleger, Stefano F.F. Torriani, Anne Dérédec, Florence Carpentier, Anne-Sophie Walker

## Abstract

Sustainable crop protection is crucial for food security, but is threatened by the adaptation of diverse, evolving pathogen population. Resistance can be managed by maximizing selection pressure diversity, by dose variation and the spatial and temporal combination of active ingredients. We explored the interplay between operational drivers for maximizing management strategy sustainability relative to the resistance status of fungal populations. We applied an experimental evolution approach to three artificial populations of an economically important wheat pathogen, *Zymoseptoria tritici,* differing in initial resistance status. We revealed that diversified selection pressure limited the selection of resistance in naïve populations and those with low frequencies of single. Increasing the number of modes of action delayed resistance development most effectively — ahead of increasing the number of fungicides, fungicide choice based on resistance risk and temporal variation in fungicide exposure — but favored generalism in the evolved populations. However, the prior presence of multiple resistant resistant isolates and their subsequent selection in populations overrode the effects of diversity in management strategies, incidentally invalidating any universal ranking. Initial resistance composition must therefore be considered specifically in sustainable resistance management, to address real-world field situation.

**Abbreviated summary:** Experimental evolution is a relevant tool for exploring the determinants of antifungal adaptation in fungi. Here, using the model species *Z. tritici* and two fungicidal modes of action associated with contrasting resistance risks, we have demonstrated that initial population composition, and in particular the presence of multiple resistance, represents the main determinant of adaptive trajectories.

**Originality-Significance Statement:** Pesticides are part of microbe environment in agricultural systems and may select for resistance. This adaptation of pathogens is a burden for plant health. Using an original multicriteria assessment and experimental evolution, we revealed that multi-fungicide use, selecting for multiple resistance, trumped any other driver of selection, invalidating any universal ranking of management strategies, a dispute extensively illustrated in literature and still inconclusive, especially in agriculture. This outcome refocuses the debate on population diversity and evolution rather than on the intrinsic properties of strategies, as mostly acclaimed in literature. From a practical point of view, our results highlight the importance of considering local population composition when recommending spraying programs. This practice is currently not generalized in agriculture but may be timely to effectively delay resistance evolution *and* reduce pesticide load in agricultural systems, a growing social demand, since resistance monitoring at relatively fine spatial scales and at low frequency may become commonplace in a near future with the rise of new molecular biology technologies

## Introduction

Pesticides or drugs exert selection pressures on microbes and pest populations, leading to the emergence of uncontrolled resistant individuals, resulting in losses of efficacy and a need for dose increases, compromising crop, animal and human health and aggravating the environmental impact of the chemicals used (McKenna, 2013; Fisher et al., 2018; Gould et al., 2018; Murray et al., 2022). Resistance management strategies aim to delay the evolution of resistance by subjecting populations to diverse and heterogeneous selection pressures or by reducing their exposure to selection (REX_Consortium, 2013; van den Bosch et al., 2014a). Selection can thus be mitigated by increasing its *diversity* (*i.e.* number of active ingredients (AIs) or modes of action (MoAs) deployed, choosing AIs according to their intrinsic risk of resistance development), or by increasing its *heterogeneity* (*i.e.* playing with pattern of application over time, either concomitantly as in mixtures, or successively *as* in alternation or cycling, or over space as in mosaic) or by varying its intensity (*i.e.* modulation of fungicide dose or number of sprays).

In agriculture, the ascomycete *Zymoseptoria tritici,* a hemibiotrophic plant pathogenic fungus responsible for Septoria tritici blotch (STB), the leading yield-reducing disease of wheat worldwide, provides an illustration of the debate about the relative performances of resistance management strategies (Fones and Gurr, 2015; Torriani et al., 2015). This pathogen is currently controlled principally by fungicide applications. This has led to its adaptation, to various degrees, to almost all the single-site MoAs authorized (benzimidazoles, which inhibit β-tubulin polymerization; QoIs, which inhibit complex III of mitochondrial respiration; DMIs, which inhibit sterol biosynthesis and SDHIs, which inhibit complex II of mitochondrial respiration), at least in Western Europe (Hellin et al., 2020; Garnault et al., 2021). Interestingly, this fungus displays various resistant phenotypes (Heick et al., 2017a; Huf et al., 2018; Jorgensen et al., 2018; Rehfus et al., 2018; Samils et al., 2021). Specialist resistance results from the selection of mutations in genes encoding the target proteins of these fungicides (target-site resistance or TSR). The combination of multiple mutations in the same isolate, resulting in multiple resistance, is favored by the high rates of sexual reproduction in this pathogen (Singh et al., 2021). Generalist resistance may result from non-target-site resistance (NTSR), as in the multidrug resistance (MDR) caused by the enhanced efflux of unrelated fungicides (Leroux and Walker, 2011; Omrane et al., 2015). The diversity of resistant phenotypes observed in *Z. tritici* and their contrasting local frequencies in the field renders this pathogen a relevant biological model for investigations of the interplay between the drivers of the performance of anti-resistance strategies .

Several empirical and modelling studies in *Zymoseptoria tritici* have concluded that strategies including alternation, mixtures, and modulations of the dose, number and timing of applications can effectively limit the evolution of resistance, provided that they are optimized (Hobbelen et al., 2013; van den Bosch et al., 2014a; Dooley et al., 2016a; Heick et al., 2017a; Jørgensen et al., 2017; Mikaberidze et al., 2017). However, it remains a matter of debate which of these strategies is the most sustainable. Attempts to rank strategies have suggested that mixture-based strategies may be the best, but the results obtained were not unequivocal for *Z. tritici* or for many agricultural or clinical situations (REX_Consortium, 2013; van den Bosch et al., 2014a). There may be several reasons for this. First, the results for strategy performance may depend on the criteria used for assessment (Elderfield et al., 2018). Indeed, the evolution or frequency of resistance are among the most widely used criteria, but many other criteria, such as disease control or fungicide effective life, may be just as relevant and may alter the ranking of performance between strategies. Furthermore, strategies must be truly comparable to obtain meaningful results. In most of the empirical data available, alternation and mixture strategies are rarely compared at similar dose of AIs, but rather at their field rates, in most instances designed not solely for fungicide resistance management but for a combination of factors, including spectrum of control, (eco)toxicological aspects and cost-effectiveness of the product. And when alternation and mixture strategies are fairly compared, the ranking is not decisive (Hobbelen et al., 2013; van den Bosch et al., 2014b). Finally, the composition of the pathogen population in terms of resistance types and frequencies (*i.e.* resistance status) may also have an impact on strategy performance, as resistant individuals are fitter than susceptible individuals in sprayed environments and higher doses of fungicide are required to control them. Furthermore, when single-resistant genotypes are already present, the acquisition of multiple resistance to two independent MoAs only requires the subsequent selection of a single new mutation. The emergence of multiple resistance is therefore more likely than when a combination of the same MoAs is used on a totally susceptible population. The drivers of heterogeneity and diversity in selection pressure may, therefore, have different impacts on resistance management, according to how and when they are implemented with respect to pathogen resistance dynamics, which depends on the current composition of the pathogen population. For example, strategies that effectively delay the initial emergence of resistance may be different from those most effective at slowing the selection of resistances already present in the pathogen population (see an example on mixture strategies (Hobbelen et al., 2014)). The initial frequency and nature of resistance mutations (*e.g.* inducing resistance to one or multiple MoAs) may also influence the effective life of AIs (Hobbelen et al., 2013). Thus, the overall performance of a strategy depends on the respective efficacies of its components for controlling both susceptible and resistant individuals over time and space, whatever the composition of the pathogen population in terms of resistance status.

In this context, we investigated the interplay between the operational drivers maximizing the sustainability of management strategies relative to the resistance status of the pathogen population. We explored several assumptions, based on published studies, including the work of Ballu *et al* (Ballu et al., 2021; Ballu et al., 2023). We found that the various sources of heterogeneity and diversity in selection pressures differed in their efficacy for delaying resistance evolution. We found that combining these sources would probably increase the overall sustainability of strategies (REX_Consortium, 2013; Ballu et al., 2023), and that the efficacy of anti-resistance strategies depends strongly on pathogen population composition in terms of resistance (Hobbelen et al., 2013). These results have been achieved through experimental evolution, an innovative approach that allows for the design of complex experiments but remains underused in fungi. *Z. tritici* is particularly suitable for investigations with this approach because of its haploid yeast-like form (*i.e.* blastospores) that is easy to culture in liquid medium. By evolving strains carrying field resistances, we were able to combine some of the characteristics of natural populations with accelerated experimental evolution in miniaturized controlled conditions (Bailey and Bataillon, 2016; Fisher and Lang, 2016). We followed the dynamics of populations and resistances under several selection regimes, which helped to elucidate the impact of population composition on the efficacy of the drivers of resistance management strategies. Several sources of diversity and heterogeneity generally considered to promote the durable control of STB were tested: the number of AIs used (*i.e.* 1-3), the number of MoAs (*i.e.* 1-3), the intrinsic vulnerability of these agents to the development of resistance (*i.e.* low to high resistance risk), and the temporal heterogeneity of fungicide exposure (*i.e.* mixture *vs.* alternation). We compared the performance of several selection strategies, defined as combinations of one or several of these drivers, against artificial populations of different compositions in terms of resistance status. We were especially interested in the introduction of multiple resistance, which is associated to the combination, in a single individual, of independent resistance determinants each involving resistance to a single mode of action, into artificial populations, to mimic some aspects of the recombination that occurs naturally in field populations of *Z. tritici.* This made it possible to explore the potential effect of such events on strategy performance in an artificial experimental design (Fisher and Lang, 2016). A multi-criteria assessment of strategies, based on growth dynamics of populations and evolution of resistance phenotype profiles, confirmed that diversification of selection pressure, by alternation or mixture-based approaches, can limit the emergence and selection of resistance, at the cost of selecting generalist or multiple resistance. But above all, resistance status of pathogen population prior to the implementation of strategies was found to be the primary determinant of strategy performance, particularly when multiple resistance was already present in the initial population. These findings open up new perspectives for the informed tailoring of strategies and show that population composition must be given priority over other drivers of selection in such strategies.

## Materials and methods

### Ancestral isolates

The susceptible ancestral isolate was the IPO-323 strain, the source of the reference genome for *Z. tritici (Goodwin et al., 2011)*, hereafter referred to as ‘S’. Any bias in fitness due to genetic background was minimized by selecting resistant isolates from the F1 progeny of a cross between ‘S’ and a resistant field strain. The SDHI-resistant isolate, RSDHI, was characterized by the substitution of an arginine (R) for the histidine (H) residue at codon 152 of the sdhC protein (*sdhC^H152R^* resistance allele), one of the subunits of succinate dehydrogenase, the target of SDHIs (Rehfus et al., 2018). The DMI-resistant isolate, RDMI, displayed five amino-acid changes (V136A, A379G, I381V, Y461S and S524T) in the sterol 14α-demethylase protein encoded by *cyp51* (the resistance allele is referred to hereafter as *cyp51^I381V^*). An isolate exhibiting multiple resistance, RSDHI/DMI, carried both unrelated resistance alleles, as well as the G143A change in cytochrome b (the target of QoI fungicides), inherited from its wild parent. The RSDHI and RDMI isolates might be referred hereafter as “single-resistant isolates”, whereas RSDHI/DMI might be referred as “the multiple-resistant isolate”.

### Fungicide sensitivity of the ancestral isolates

We established the resistance profile of each strain by estimating EC50 values and resistance factors (RFs) for a wide range of fungicides. The fungicides studied were all of technical grade and included three SDHIs (benzovindiflupyr, boscalid and fluxapyroxad), three DMIs (prothioconazole-desthio, metconazole and mefentrifluconazole), a QoI (azoxystrobin), a benzimidazole (carbendazim), a multisite inhibitor (chlorothalonil) and tolnaftate (an inhibitor of squalene epoxidase, an enzyme involved in sterol biosynthesis, used in medicine and as a marker of multidrug resistance). Fungicides were dissolved in DMSO and stock solutions were prepared so as to ensure that the final medium contained no more than 0.5% DMSO.

EC50 values were calculated from dose-response curves plotted for data obtained in 96-well microtiter plates (Sarstedt, Germany). Pure liquid cultures of each isolate were grown in 50 mL borosilicate Erlenmeyer flasks containing 25 mL YPD medium (20 g.L^-1^ dextrose, 20 g.L^-1^ peptone, 10 g.L^-1^ yeast extract; USBiological Salem, MA, USA), with carded cotton wool inserted into the neck. The cultures were maintained at 18°C and a RH of 70% in the dark for 7 days, with shaking at 150 rpm. These cultures were used to inoculate 200 µL of YPD medium per well in microtiter plates, at an initial density of 2.5 x 10^5^ spores.mL^-1^. We tested 10 fungicide concentrations for each fungicide, in addition to a control amended only with 0.5% DMSO. The design included three replicates per isolate and per fungicide concentration. The plate was sealed with a gas-permeable membrane (Breathe Easy®; Diversified Biotech) and placed on a shaker in the dark, in the conditions described above. Optical density at 405 nm (OD405) (SpectraMax M2 spectrophotometer, Molecular Devices) was measured after five days, *i.e.* at the end of the exponential growth phase, with three technical replicates of each well. Each experiment was repeated at least three times.

EC50 values were determined by logistic regression (logistic nonlinear mixed effects model, with the SSlogis function of the Self-Starting Nls Logistic Model, available in the STATS package and R version 4.1.3), with dose as a fixed effect and biological replicate as a random effect.

RFs were calculated as the ratio of the EC50 of the resistant isolate to that of the susceptible isolate. Standard errors were calculated as follows:

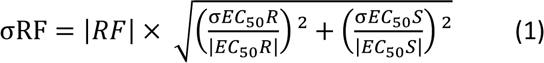

where R is the resistant isolate and S is the susceptible isolate.

Fitness and resistance profiles of ancestral strains

Fitness was inferred for each isolate from its growth in the culture conditions used for experimental evolution (detailed below) but without fungicides. Population size was measured daily on the basis of OD405 data, for seven days, as previously described (Ballu et al., 2021). The experiment was performed five times. The growth curve for each strain was modeled with a mixed non-linear logistic model, as follows:

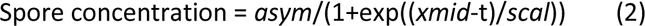

where *t* is the day, *asym* the asymptote, *i.e.* maximum growth, *xmid* is the time required to decrease the value of the asymptote by half and *scal* is the inverse of the slope.

The strains were considered as a fixed effect, with replicate as a random effect. We compared parameters between strains with the Tukey method for pairwise comparisons. Estimates were obtained with the NLME, CAR and EMMEANS packages of R version 4.1.3.

### Artificial ancestral populations

Three ancestral populations were prepared for the first round of experimental evolution by inoculating medium with pure cultures of ancestral isolates (grown as described in the previous section) in various proportions so as to obtain a final density of 10^7^ spores in total in 25 mL YPD medium amended with 0.25% antibiotics (100 mg.L^-1^ streptomycin and penicillin – to prevent contamination in this long-term experiment) in 50 mL Erlenmeyer flasks (**Fig. 1C**). PopS was a naïve population composed exclusively of the susceptible isolate S. PopR was composed of 95% S and of 5% RDMI and RSDHI (1:1) and therefore contained only single-resistance isolates although both resistances coexisted in the same population. PopRR contained 95% S and 5% RDMI, RSDHI and RSDHI/DMI (2:2:1) and therefore contained a collection of single- and multiple-resistant isolates. This initial population composition of PopR and PopRR was determined in preliminary experiments: a proportion of 1% was found to be the smallest proportion providing the lowest mean and standard deviation of experimental error in resistance allele quantification by qPCR or pyrosequencing (see below) on a range of dilutions. As multiple resistance was expected to be scarce in natural populations, we arbitrarily set the initial frequency of RSDHI/DMI at 1% and half that of single-resistant isolates.

**Figure 1:**
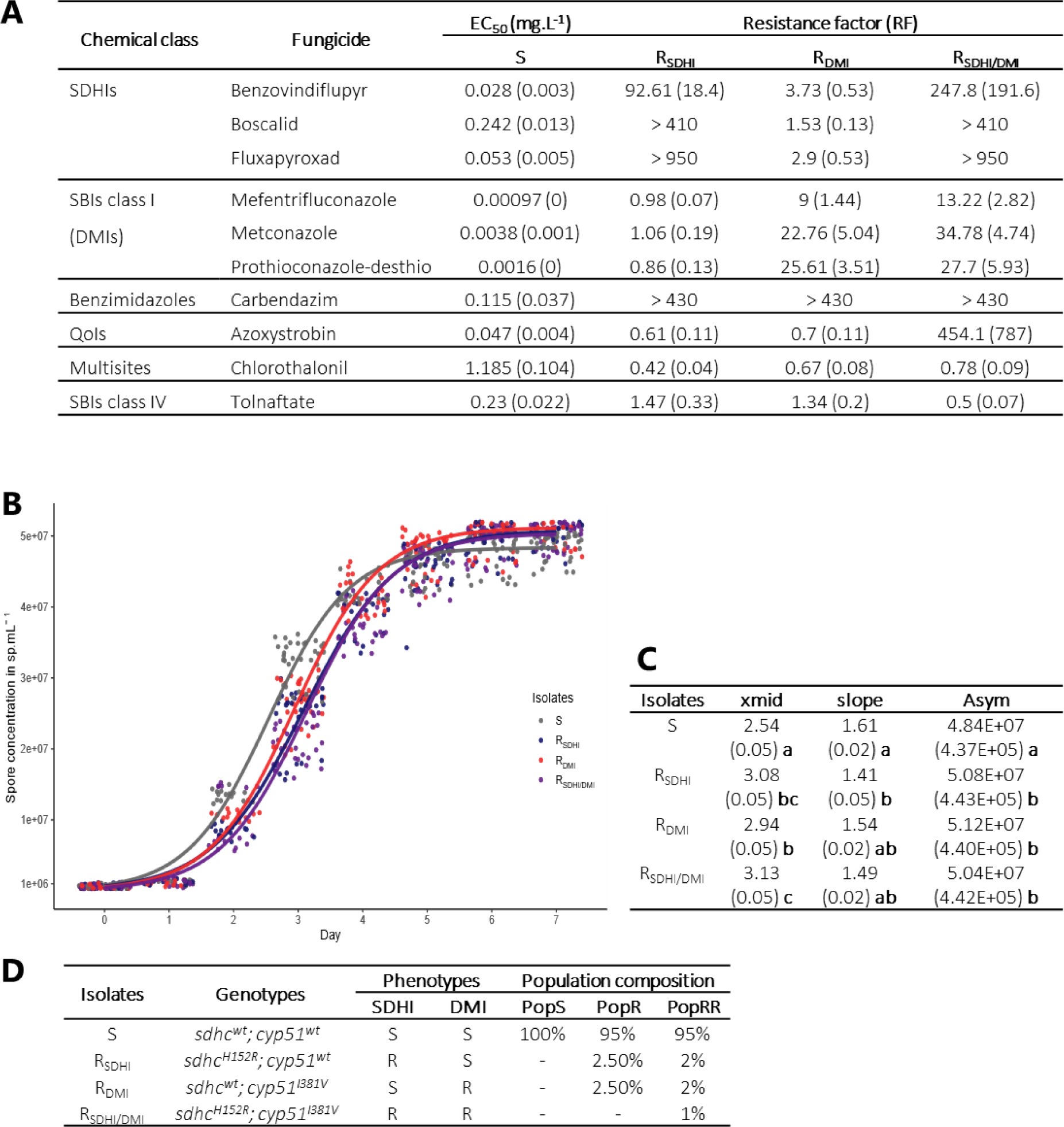
***Features of the ancestral isolates and populations of Z. tritici. used in experimental evolution.*** The S, RSDHI, RDMI and RSDHI/DMI isolates are the ancestral isolates used in the experiment fully susceptible, resistant to SDHIs, resistant to DMIs and resistant to both, respectively (Dooley et al., 2016b; Huf et al., 2018). ***A. Profiles of susceptibility to multiple fungicides of ancestral isolates of* Z. tritici.** Estimations of EC50 and Resistance Factors (*i.e.* EC50 R isolate / EC50 S isolate). The standard error is specified in brackets. ***B. Fitness of ancestral isolates in the conditions used for experimental evolution.*** Growth without fungicide, for cultures of single isolates. The experiment was performed five times and the data were modeled with a mixed non-linear logistic regression model. ***C. Estimated parameters of the growth curves for ancestral isolates.*** The standard error is shown in brackets. *asym* is the asymptote (*i.e.* maximum growth); *xmid* is the time required to decrease the asymptote by 50%; *scal is* the inverse of the slope*. The group letters refer to Tukey’s pairwise tests; different letters indicate significant differences (P<0.05). **D. Artificial populations of*** **Z. tritici *used for experimental evolution.*** The evolution of a fully susceptible PopS population was compared to that of the PopR and PopRR populations containing only single-resistant isolates and single- and multiple-resistant isolates, respectively.

### Components of selection regimes diversity

The 12 selection regimes differed in terms of four components and are described in detail in **Supplementary Information 2**.

The number of AIs ranged from one (straight selection) to three (diversified selection).

The number of MoAs ranged from one (intra-MoA selection) to three (inter-MoA selection).

We selected AIs representative of their MoAs from among those currently used to control STB in the field. MoAs, and, thus, their representative AIs, differed in terms of their vulnerability to resistance, *i.e.* intrinsic risk of resistance (Brent and Hollomon, 2007). In our view, the risk of resistance is seen globally and may include the time to resistance emergence, the speed of resistance selection and resistance intensity and pattern, as determined by the specific resistance mechanism(s) selected. Selected AIs were considered to be at medium-to-high risk (SDHIs, represented by benzovindiflupyr (B) and fluxapyroxad (F)), low-to-medium risk (DMIs, represented by prothioconazole-desthio (P), the active metabolite of the commercialized prothioconazole, and mefentrifluconazole (M)) or at very low risk (multisite inhibitor, represented by chlorothalonil (C)) of resistance.

We also studied the effect of temporal heterogeneity through comparisons of the simultaneous application of AIs (intracycle heterogeneity, *i.e.* mixture) with the sequential application of AIs (intercycle heterogeneity, *i.e.* alternation). All the alternation regimes began with the AI with the lowest risk of resistance (or highest intrinsic activity for strategies with only one MoA), as this has been shown to be the most effective approach (van den Bosch and Gilligan, 2008; Elderfield et al., 2018).

### Regimes of selection applied to artificial populations

These 12 selection regimes and 2 control situations were applied to all or some of the three possible ancestral populations, resulting in 31 possible interactions, and 100 independent lines, as there were three replicates for each interaction (or four when chlorothalonil was involved) and five solvent-only controls were included (**Supplementary Information 2)**. Given the large size of this experiment, selection regimes were grouped together into two batches (A and B) performed successively. Strategies that it was particularly important to compare directly were included in the same batch. The two control lines included only 0.5% of DMSO or ethanol 80%. Controls were repeated in both batches if necessary.

### Fungicides preparation

Fungicides were used as technical products purchased from Sigma Aldrich (US). They were dissolved in 80% ethanol, with the exception of chlorothalonil, which was dissolved in DMSO due to its poor solubility in ethanol. Controls in which only solvent (80% ethanol or DMSO) was added were included in the experiment. Stock solutions of fungicides were prepared at the beginning of the experiment and were stored at 4°C. The fungicides and solvents added to culture medium never exceeded 0.5% of the final volume, to prevent toxic effects on the growth of *Z. tritici*.

### Selection doses

For fungicides used alone (in sequence or in alternation), the selection dose was the EC97 of each AI determined in a preliminary experiment for the S isolate grown in the conditions used for experimental evolution (described below). EC97 values were established from dose-response curves for each fungicide based on the logistic regression model used to describe the resistance patterns of the ancestral isolates. The growth of the populations, inferred from OD405 measurement after 7 days of culture, was normalized by dividing by that of the control line grown without fungicide. The validation of this selection dose included culture of the S isolate in the conditions used for experimental evolution at its theoretical EC97 and at close doses. The lowest dose providing at least 97% inhibition was finally defined as the selection dose.

The selection doses of two-way fungicide mixtures were set at half the selection dose of each AI used alone, for the comparison of alternation and mixture regimes containing the same amounts of each fungicide. The only three-way mixture studied contained half the EC97 of chlorothalonil, a quarter the EC97 of benzovindiflupyr and a quarter the EC97 of prothioconazole-desthio.

The selection doses are detailed in **Supplementary Information 2**.

### Design of the experimental evolution experiment

Experimental evolution was performed over either six or eight cycles of seven days each (about six to seven generations per cycle), depending on the experimental batch. This cycle duration corresponds to about the time for which a control line remained in exponential growth phase before reaching a plateau. On the first day of the experiment, ancestral population lines containing 10^7^ spores were prepared and incubated as described above, after amendment with the fungicides required for each selection regime. OD405 was measured at the end of each cycle and transformed into population size, as established in preliminary experiments (See more details in Ballu et al., 2023). The OD405 of the treated lines was normalized against the mean OD405 for the control solvent lines. At each transfer, 500 µL of the evolving culture (*i.e.* 2% of the total volume of medium) or a maximum of 10^7^ spores was transferred to a new Erlenmeyer flask containing fresh medium. If the number of spores in the 500 µL of culture medium was less than 10^7^, as would be the case before the evolution of resistance, then the appropriate number of spores from the appropriate source population (*i.e.* spores from the control (PopS) or mixtures of spores reconstituting PopR or PopRR from the pure cultures of the different isolates grown separately without fungicides) was added to make the number of spores present up to 10^7^. Thereby the starting population size was equalized between lines at the start of each new cycle, and the process mimicked the immigration occurring in field situations. For each of the three replicates, a dedicated source population was used for immigration throughout the experiment. In preliminary experiments, immigration was found to be useful for preventing population extinction after weekly bottlenecks, and to minimize genetic drift.

At the end of each cycle, 2 mL of each line was mixed with glycerol (25%), frozen and stored at -80°C for further analyses. If the normalized Malthusian growth (defined below) of a line reached at least 95% during three consecutives cycles, the line was ended. In such cases, we ensured fair comparison between alternation and mixture approaches by stopping lines only after an even number of cycles (*i.e.* not before the 4^th^ cycle). In these situations, data (population size or allele content) for the missing cycles during this establishment phase were extrapolated for analyses involving comparisons between selection regimes. Extrapolated data were calculated using the average value of the last three consecutive cyles with generalized resistance normalized by divided by the value of the control at the considered cycle.

### Assessment of the rate of resistance evolution

The Malthusian growth (Fisher, 1999) of each line was calculated as

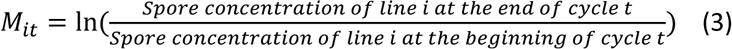

and normalized against that of the solvent control line

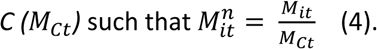

For each selection regime, the mean Malthusian growth *M*^*n*^_*i*_ is the mean of *M*^*n*^_*it*_ calculated over the total number of cycles the line was grown, and constitutes a quantitative summary of the increase in resistance of the three replicates over the entire experiment.

We used beta regression (package BETAREG for R) on the *M*^*n*^_*i*_ of each line to investigate the effects of population composition and of the components of selection diversity and temporal heterogeneity, included as fixed factors, with experimental batch as a random factor. These models are detailed in **Supplementary Information 7**. We evaluated the impact of these factors with an analysis-of-variance-like (joint_tests function of EMMEANS package) analysis, with the Tukey adjustment method for pairwise comparisons (package EMMEANS).

Molecular quantification of resistance alleles in evolved lines

The *sdhC^H152R^* allele, included in the RSDHI and RSDHI/DMI ancestral isolates, was quantified by qPCR. The *cyp51^I381V^* allele, included in the RDMI and RSDHI/DMI ancestral isolates, was quantified by pyrosequencing. The *cytb^G143A^* allele, included only in the RSDHI/DMI ancestral isolate was quantified by qPCR. Details on these procedures are provided as **Supplementary Information 4**.

From qPCR and pyrosequencing data, we calculated *RC*^*n*^_*it*_, the normalized concentration of the resistant mutants RSDHI and RDMI in the population, as the proportion of the resistance alleles *sdhC^H152R^* and *cyp51^I381V^*, respectively, multiplied by the size of the population (in spores.mL^-1^):

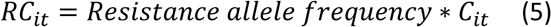

where *C*_*it*_ is the concentration of the line *i* at time *t*.

We then divided this value by the spore concentration of the control lines, *C*_*ct*_, to obtain the normalized concentration of resistance (hereafter referred to as the normalized allelic concentration):

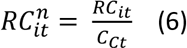

The qPCR and pyrosequencing methods used could not distinguish combinations of *sdhC^H152R^* and *cyp51^I381V^* within the same individual (*i.e.* as in RSDHI/DMI) from co-occurrence of the same alleles in different individuals (*i.e.* as in the RSDHI and RDMI isolates). We therefore set the maximum possible concentration of this multiple-resistant isolate as the lower proportion of the two resistant alleles compared together. This calculation may overestimate the concentration of the RSDHI/DMI isolate in PopRR. For validation of this estimate, we compared these data with those for *cytb^G143A^* quantification on samples collected from the last cycles of PopRR lines.

For each of the three resistance alleles, we used beta regression (package BETAREG for R) on the mean normalized allele concentrations of each line to investigate the effects of population composition, with the components of selection diversity and temporal heterogeneity treated as fixed factors, and experimental batch as a random factor. We evaluated the impact of factors with an analysis-of-variance-like (joint_tests function of EMMEANS package) approach with the Tukey adjustment method for pairwise comparisons (package EMMEANS). Details of these models are provided in **Supplementary Information 7** and models are referred hereafter as MOD”*NAME”*.

### Phenotype resistance profiles of evolved lines

The cross-resistance patterns of all populations collected at the end of the experimental evolution were established in droplet tests. We deposited population suspensions containing 10^7^, 10^6^, 10^5^, and 10^4^ spores.mL^-1^ as drops organized into columns, on 12 cm square Petri dishes. These plates contained YPD supplemented with discriminatory doses of fungicides, as detailed in **Supplementary Information 3**. The 10 treatments tested encompassed a control with 0.5% solvent and fungicides B, P, F, M and C at their selection doses and their mixtures. Finally, for the detection of generalist resistance, we also included fungicides not used for experimental evolution (tolnaftate and azoxystrobin although this fungicide is of limited interest for PopRR lines where the *cytb^G143A^* was introduced with the RSDHI/DMI isolate). Tolnaftate, in particular, identifies individuals displaying enhanced fungicide efflux leading to non-specific multidrug resistance (MDR), as described in field isolates of *Z. tritici* (Leroux and Walker, 2011). Ancestral isolates, and a MDR isolate collected in the field were included as control strains. After seven days of incubation, drops were scored 0 or 1, according to the presence or absence of fungal growth (**Fig. 5A**). A total score was established for each strain, ranging from 0 (susceptible) to 4 (resistant if >0), according to the number of dilution droplets for which growth was observed. For each strain, the resistance phenotype profile was determined by concatenating the scores established for the 10 sets of fungicide conditions.

The distributions of resistance phenotype profiles between and within selection regimes were plotted as heatmaps. The Euclidean pairwise distance was used for the hierarchical clustering of these profiles, with dendrograms for the rows and columns, in R version 4.1.3. A principal component analysis (PCA) was also performed on the same phenotypes.

### Characterization of enhanced efflux

The 17 strains isolated from lines growing on tolnaftate in droplet tests were selected for further investigation. We checked for changes in the length of the promoter of *mfs1* by PCR, as previously described (Omrane et al., 2017). Several types of insertion into the promoter of this gene encoding a MFS transporter have been shown to cause enhanced efflux leading to multidrug resistance in field isolates of *Z. tritici*. We quantified *mfs1* expression in the same isolates by RT-qPCR (Omrane et al., 2015). The protocols used are detailed in **Supplementary Information 6**.

## Results

### Ancestral isolates display contrasting patterns of resistance in artificial populations

Experimental evolution was performed with artificial populations constituted from ancestral isolates exhibiting resistance alleles currently found in European populations of *Z. tritici*. The resistant isolates (RSDHI, RDMI and RSDHI/DMI) were isolated from a second-generation progeny of a cross between the susceptible (S) wild-type reference isolate and a field isolate bearing the *sdhC^H152R^* change and several *cyp51* substitutions (*i.e.* V136A, A379G, I381V, Y461S and S524T; hereafter simplified as *cyp51^I381V^*). We checked that the resulting changes to the target proteins of SDHI and DMI fungicides, respectively, were associated with the expected phenotypes and calculated their resistance factors (RFs) relative to the S isolate (**Fig. 1A**; **Supplementary Information 1**). We confirmed that the RSDHI and RSDHI/DMI isolates bearing *sdhC^H152R^* had RFs exceeding 90 for all SDHIs (Dooley et al., 2016b; Yamashita and Fraaije, 2018) and that the RDMI and RSDHI/DMI isolates bearing *cyp51^I381V^* displayed moderate resistance to all DMIs tested (9<RFs<40), although the RFs for mefentrifluconazole were lower than those for other azoles (Huf et al., 2018; Ishii et al., 2020). Complementary genotyping revealed that the RSDHI, RDMI and RSDHI/DMI isolates had inherited resistance to benzimidazoles from their parental field isolate, precluding any further use of these fungicides in selection regimes (Hawkins and Fraaije, 2016), and that only the multiple-resistant RSDHI/DMI isolate had inherited resistance to QoIs due to the *cytb^G143A^*change (Torriani et al., 2015). All isolates were equally susceptible to the multisite agent chlorothalonil and to tolnaftate, an inhibitor of squalene epoxidase used to reveal multidrug resistance in *Z. tritici* (Leroux and Walker, 2011).

The fitness of the ancestral isolates was assessed by establishing growth curves under the conditions used for experimental evolution and in the absence of fungicide selection (**Fig. 1B**), and the parameters of the logistic model were used to compare isolates (**Fig. 1C**). The time required to reach the half-maximal concentration of spores was significantly greater for the resistant isolates than for the reference isolate (2.94-3.08 days *vs.* 2.5 days), but maximal spore concentration was significantly higher for resistant isolates than for the reference isolate at the end of the experiment (+4.1 - +5.8%). However, in populations from control lines not exposed to fungicide (see below), the resistance alleles initially present at low frequency were not detected at the end of the experimental evolution period, suggesting that either the weak and early penalty associated with resistance observed in our conditions was expressed, and/or that they were lost by chance because of the regular bottlenecks affecting populations during their serial cultivation. As no other genotype replicates were tested, it remains unclear whether this fitness penalty was related to the resistance mutations or to the genetic background of the isolates. Nevertheless, these findings clarify and complete previous observations concerning the fitness cost associated with the *sdhC^H152R^* variant expressed in culture *in vitro* and in competition experiments (Scalliet et al., 2012; Gutiérrez-Alonso et al., 2017), and restoration of the fitness cost of the I381V variant when associated with a modification to the Y461 codon of *cyp51*, as in RDMI and RSDHI/DMI(Hawkins and Fraaije, 2018). We assumed that resistance to benzimidazoles did not bias fitness measurement, as no associated fitness penalty has ever been observed for this resistance in any species, including *Z. tritici* (Hawkins and Fraaije, 2016; Hawkins and Fraaije, 2018). Resistance to QoIs has been reported to decrease *Z. tritici* virulence *in planta* (Hawkins and Fraaije, 2018), but we assumed that this cost did not operate *in vitro* for the RSDHI/DMI isolate.

### Population composition affects the performance of resistance management strategies

We investigated the effect of population composition on the performance of strategies, by comparing the evolution of three artificial populations containing 0% (PopS) or 5% (PopR and PopRR) mixed resistant individuals, to reproduce the late stages of emergence of resistance as usually detected with current monitoring tools in field situations (**Fig. 1D**). PopR contained only single-resistant isolates (*i.e.* RSDHI and RDMI) and PopRR contained both single-resistant isolates and a small proportion of multiple-resistant individuals (*i.e.* RSDHI/DMI), to mimic the results of the sexual reproduction likely to occur in field populations of *Z. tritici* (Singh et al., 2021). These populations were subjected to selection pressure from alternating treatments with benzovindiflupyr (a SDHI; hereafter referred to as B) and prothioconazole-desthio (a DMI; hereafter referred to as P) or a mixture of these treatments, and were then compared with untreated lines (**Fig. 2A**; **Fig. 2B**; **Fig. 2C**; **Supplementary Information 2**). We quantified the dynamics of resistance in each line as its mean normalized Malthusian growth, *M*^*n*^_*i*_, a proxy for the increase in resistance under selection (**Fig. 3A**). The growth of PopS was significantly smaller (82% and 84% smaller, under P and B mixture or alternation, respectively) than that of the control. Significant decreases in growth relative to the control were also observed for the PopR population treated with the P+B mixture (85%), and to a lesser extent, the PopR population subjected to an alternation of P and B (46%). By contrast, the growth of PopRR was much more similar to that of the control population, despite very slight, but nevertheless significant decreases of 3% in alternation and 5% in mixture, revealing that both these strategies performed very poorly once multiple-resistant isolates were present in the population. These observations mirrored the strong and significant selection (increase by a factor of 53 to 157) of the resistant *sdhC^H152R^* and *cyp51^I381V^* alleles in all PopRR lines (**Fig. 3B** and **3C**), and, more specifically, of the multiple-resistant isolate RSDHI/DMI, as revealed directly by the peak frequencies of *cytb^G143A^* at the end of the experiment (**Supplementary Information 5**). The more moderate increase (increase by a factor of 0 to 12) in the frequencies of *sdhC^H152R^* and *cyp51^I381V^* in PopR lines confirmed that the various strategies, including mixture in particular, were useful for mitigating the selection of single-resistant isolates, at least during the eight cycles of the experiment (*i.e.* ≈ 50-55 generations).

**Figure 2:**
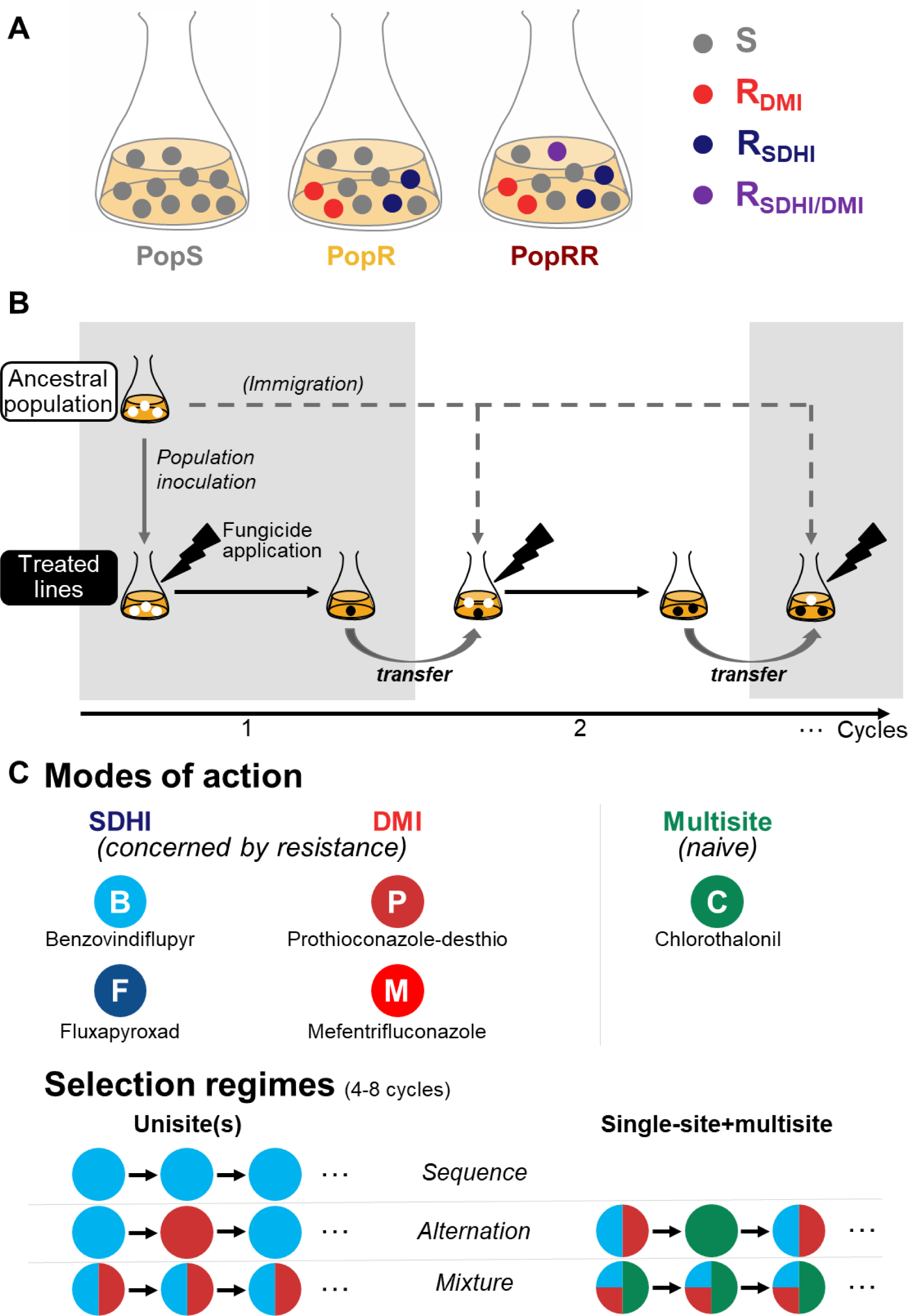
**Experimental evolution design. *A. Ancestral populations.*** Isolate S (susceptible) in gray; RSDHI (resistant to SDHIs) in blue; RDMI (resistant to DMIs) in red; RSDHI/DMI (resistant to SDHIs and DMIs) in purple. ***B. General design of the experimental evolution.*** The three ancestral populations were used to found 10^7^ spores lines. Fungicides were added to the treated lines to mimic distinct patterns of selection. After seven days, 2% of the population was used to inoculate fresh medium to start the next cycle, supplemented, if necessary, by immigration from the untreated control line from the same cycle to achieve a total of 10^7^ spores. ***C. Fungicides and selection regimes.*** Five fungicides (indicated by specific labels and colors) were studied: two SDHIs, benzovindiflupyr and fluxapyroxad (B and F; shades of blue; moderate resistance risk); two DMIs, prothioconazole-desthio and mefentrifluconazole (P and M; shades of red; low resistance risk); and the multisite inhibitor chlorothalonil (C; green; no risk of resistance). Fungicides were used in sequence, alternation or in mixture (see **Supplementary Information 2** for detailed description of selection regimes).

**Figure 3:**
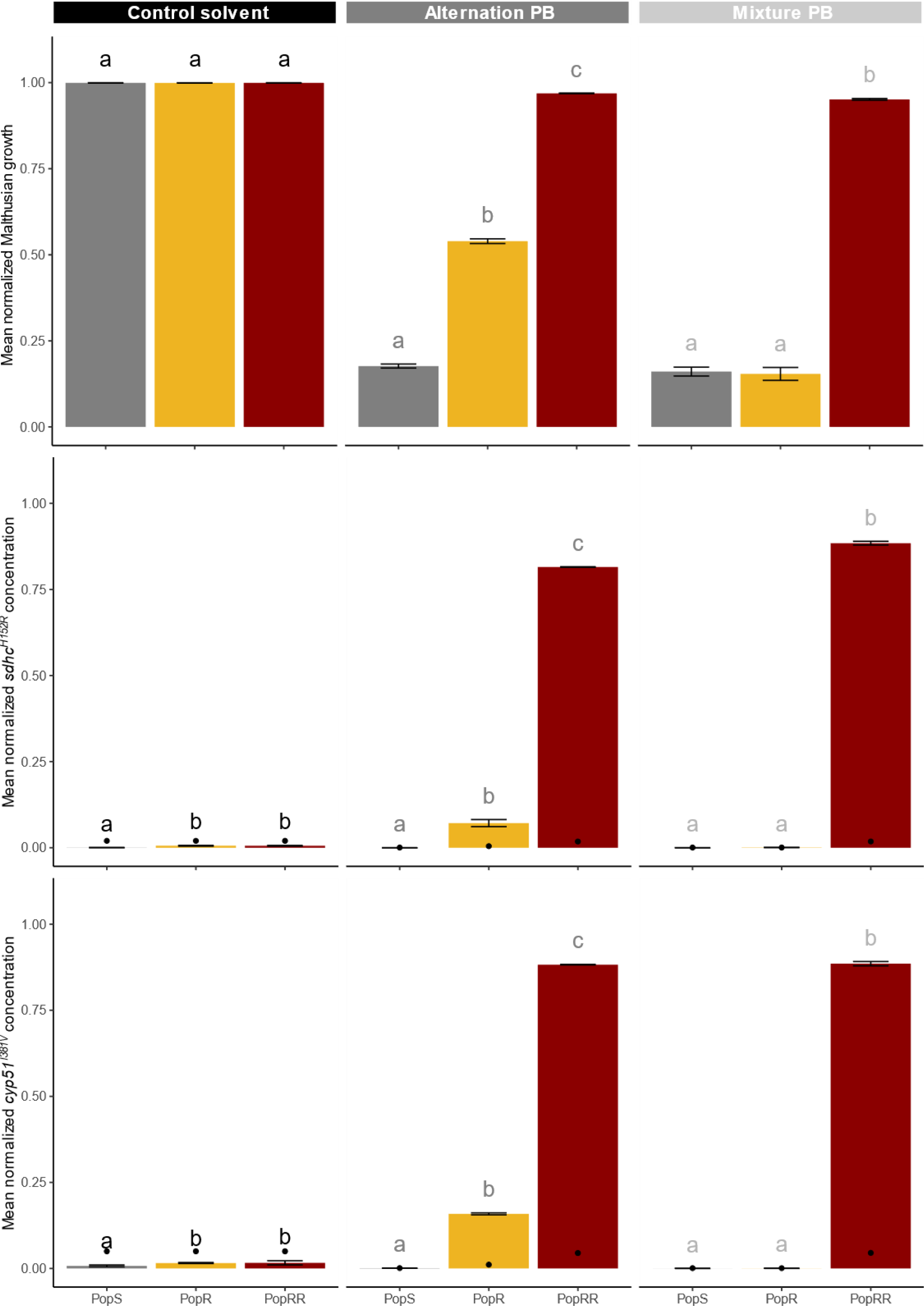
Performance of management strategies according to the resistance composition of the ancestral population of Z. *tritici*. Selection regimes included the mixture or alternation of benzovindiflupyr (B) and prothio- desthio (P), contrasted with a control amended only with 0.5% solvent. Ancestral populations (PopS, PopR and PopRR,) respectively in gray, yellow and dark red. The group letters refer to Tukey’s pairwise tests; different letters indicate significant differences (P<0.05). ***A: Mean normalized Malthusian growth. B: Mean normalized concentration of sdhc^H152R^.*** This allele confers high-level resistance to B and was quantified by qPCR. Black dots indicate the detection limit of this allele. ***C. Mean normalized concentration of cyp51^I381V^.*** This allele confers moderate resistance to P and was quantified by pyrosequencing. Black dots represent the detection limit of this allele.

We also established phenotype resistance profiles for all the lines at the end of the experiment, to observe the selection of *de novo* resistances other than those introduced *via* the ancestral phenotypes. Principal component analysis confirmed that the evolved lines differed mainly in their ancestral composition rather than in selection regimes (**Fig. 5B**). The PopS and PopRR lines retained their initial phenotypes, but one PopR line evolved resistance to tolnaftate, an inhibitor of squalene epoxidase often used to reveal multidrug resistance(Leroux and Walker, 2011).

### Mode-of-action diversity is a driver of strategy performance that should be considered before increasing the number of AIs

We investigated the extent to which population composition interfered with the components of strategies and determined their performance, by subjecting PopR and PopRR to 12 selection regimes mimicking various degrees of selection heterogeneity and diversity, and comparing them with untreated controls. We used two SDHI fungicides (B and F) potentially selecting for *sdhC^H152R^*, and two DMI fungicides (P and M) potentially selecting for *cyp51^I381V^* to investigate the effect of the number of AIs (1 *vs.* 2) and of the number of MoAs (1 *vs.* 2) on resistance evolution (**Fig. 2C** and **Supplementary Information 2**). In PopRR lines, the maximal concentration of the multiple-resistant isolate was estimated to be the lower of the concentrations of the two alleles. In situations in which *cytb^G143A^* was quantified, these two measurements were found to be highly correlated (Pearson’s *r* = 0.998; *P*<0.0001).

In strategies including two AIs with the same MoA, resistance was rapidly established, as shown by the maximal Malthusian growth and the increase in resistance allele frequencies, as early as the second cycle, as with the continuous use of a single AI, whatever the ancestral population (**Fig. 6A and Supplementary Information 8**). Such strategies even significantly accelerated the selection of resistant alleles in PopR lines relative to the continuous use of a single AI (+17.8% for *sdhC^H152R^* on average in strategies combining two SDHIs; +2.3% for *cyp51^I381V^* on average in strategies combining two DMIs). The selection of single-resistant isolates was also favored over that of multiple-resistant ones in PopRR lines (a mean allele frequency of ≈97% and ≈86% for *cyp51^I381V^* and *sdhC^H152R^*, respectively, over the experiment, *vs.* a mean frequency of 42% for multiple-resistant individuals) (**Fig. 4**, **Fig. 6A** and **Supplementary Information 5** and **8**). Moreover, the relative ratio of single- *vs.* multiple-resistant isolates for each MoA was 2:1 at the beginning of the experiment. This ratio greatly increased under DMI selection but stayed roughly stable under SDHI selection, at the end of the experiment. This finding suggests a relative resistance cost for the RSDHI/DMI isolate in competition with RDMI isolate under DMI selection pressure although both co-occurred.

**Figure 4:**
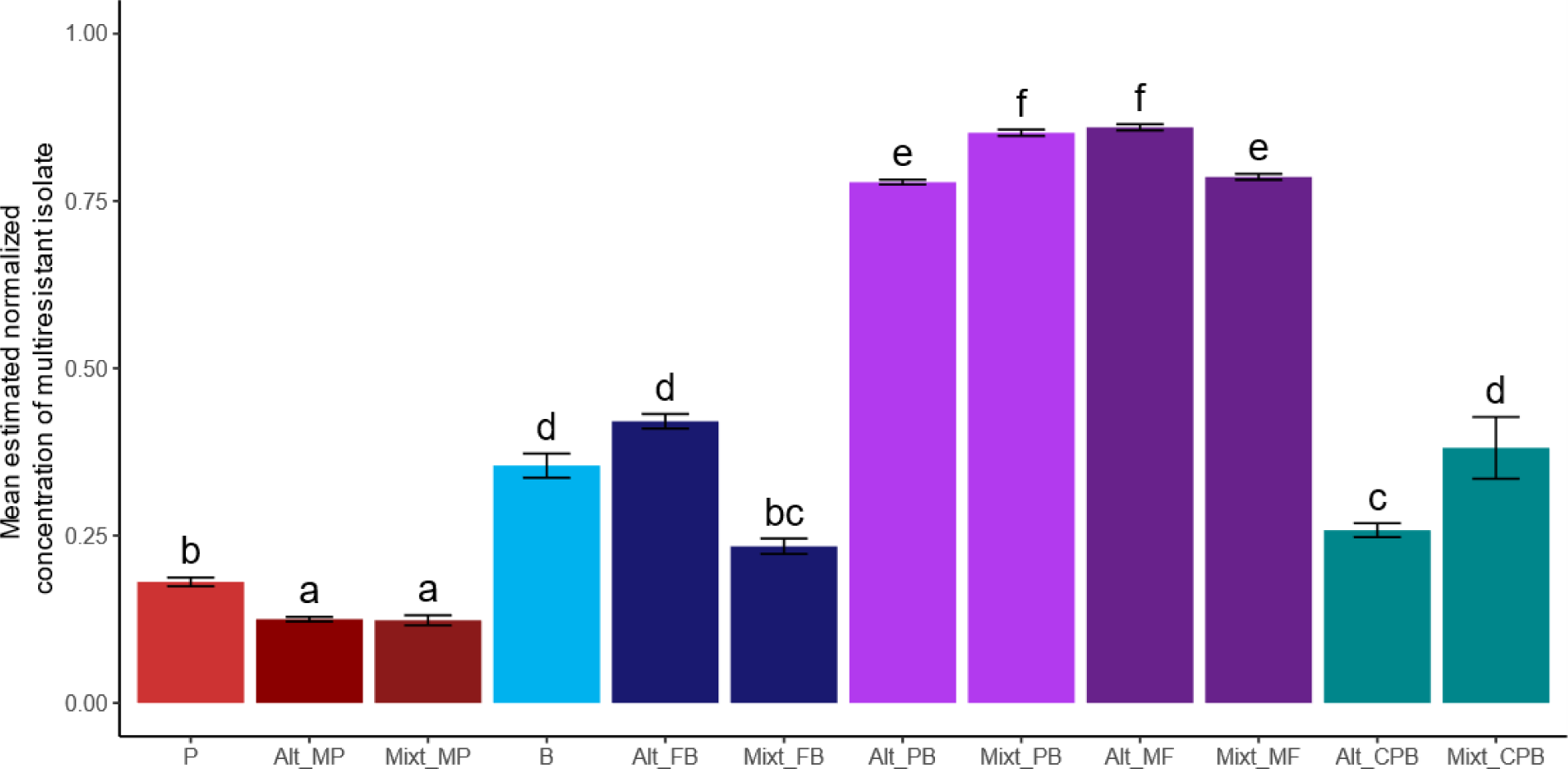
Mean maximal concentration of the multiresistant isolate in lines evolved from the PopRR ancestral population. Molecular methods used to quantify allele content could not distinguish combinations of sdhC*^H152R^ and cyp51^I381V^* within the same individual (i.e. as in RSDHI/DMI) from co-occurrence of the same alleles in different individuals (i.e. as in the RSDHI and RDMI isolates). Group letters refer to Tukey’s pairwise tests; different letters indicate significant differences (P<0.05).

**Figure 5:**
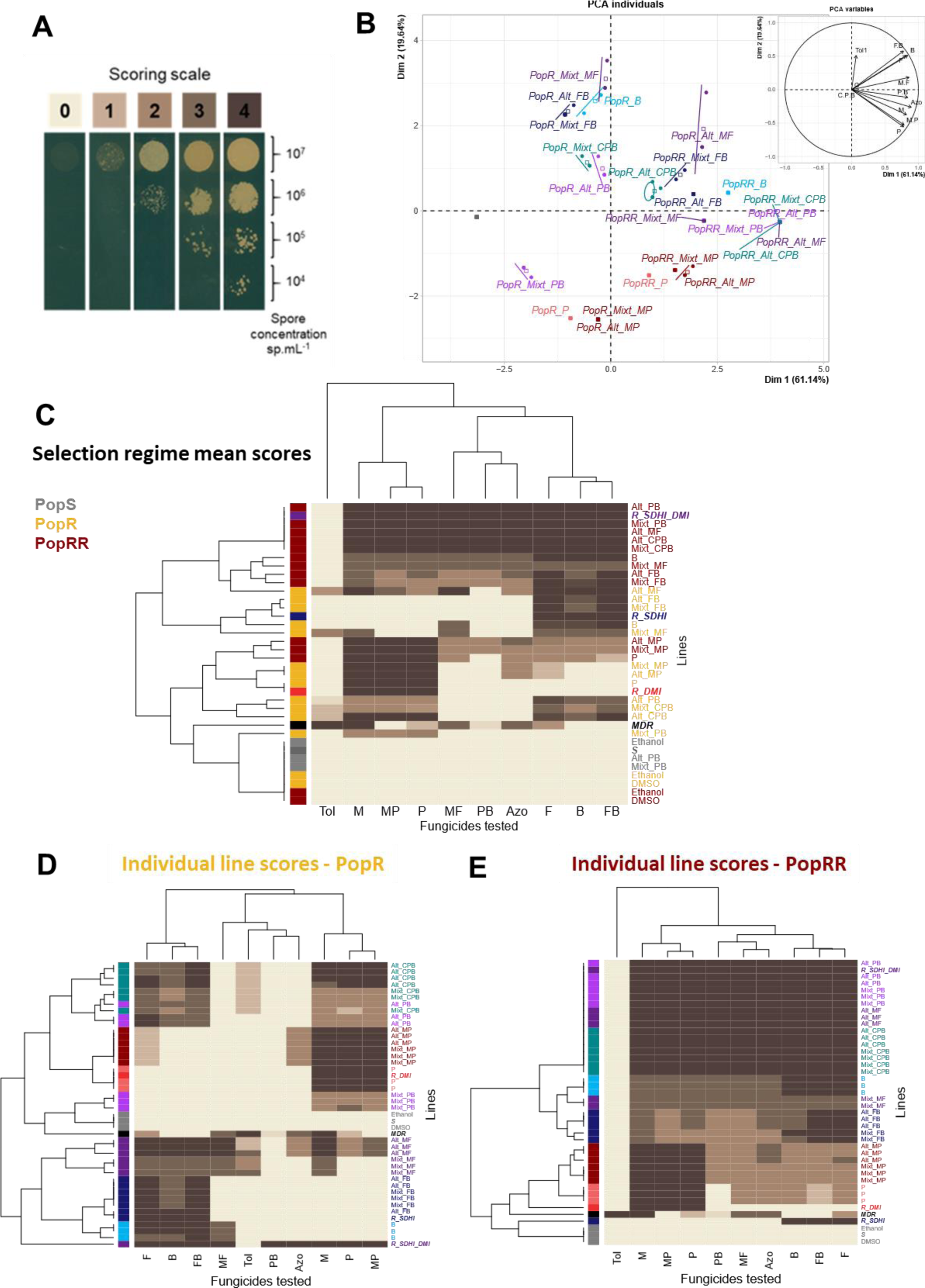
Phenotype resistance profiles of *Z. tritici* observed at the end of experimental evolution. The names of the ancestral isolates and selection regimes are as in Fig. 1 and **Supplementary Information 2**. The fungicides tested in droplet tests included benzovindiflupyr (B), fluxapyroxad (F), prothioconazole-desthio (P) and mefentrifluconazole (M) at their selection doses (EC97) and in mixture (1/2 EC97 of each), as well as azoxystrobin (Azo) and tolnaftate (Tol) which were not used for experimental evolution, at the minimal dose controlling the ancestral susceptible ancestral isolate. MDR: MDR isolate collected in the field (Leroux and Walker, 2011) serving as a reference for generalist resistance and exhibiting high resistant to tolnaftate. Scoring after the 4^th^ or 6^th^ cycle, according to selection regime. ***A. Scoring scale used to assess the growth of lines on solid medium supplemented with discriminatory doses of fungicides (droplet tests).*** For each population, four spore concentrations were tested on the same plate. 0: no growth (susceptible isolate). 1-4: growth varying with spore concentration (resistant isolate). ***B: Individual graph from the principal component analysis performed on the phenotype resistance profiles of the evolved lines***. Colors according to selection regimes (ellipses for each ancestral population line x selection regime). The inset graphic shows the PCA variables. The first axis of the PCA discriminated the evolved lines according to their ancestral populations, based on their resistance phenotype profiles. ***C: Heatmaps of average resistance phenotype profile scores for all selection regimes and ancestral populations.*** Mean of the scores established for the three population replicates subjected to the same selection regime. ***Heatmaps of phenotype resistance profile scores for individual PopR lines (D) and individual PopRR lines (E)***. Ancestral or field strains are shown in bold italics. Lines show individual scores unless no or few variability between the three repeats justifies showing their mean instead.

**Figure 6:**
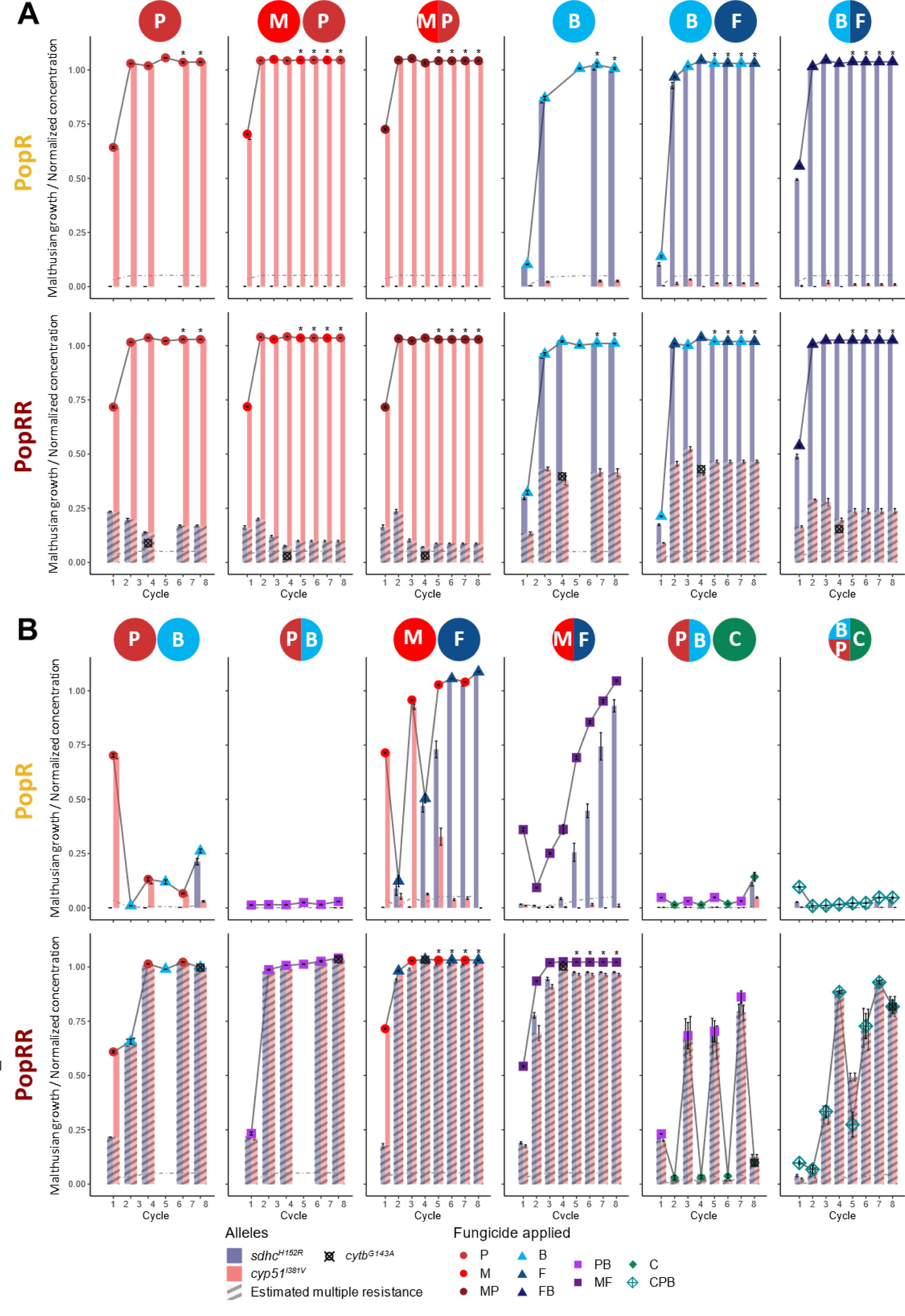
Performance of management strategies according to the selection regime applied to the ancestral population of *Z. tritici*. Selection regimes are indicated at the top of the panels and refer to the sequence (one fungicide dot), alternation (two successive fungicide dots) or mixture (one shared dot) of fungicides applied. The color and shape of the symbols in the graphs indicate the selecting fungicide used at a given cycle. Light gray lines show the change in Malthusian growth in evolved lines over the eight cycles of experimental evolution. Bars show the change in the concentration of resistance alleles (*sdhC^H152R^* in blue and *cyp5^I381V^* in red). The dashed portion of these bars corresponds to the maximum possible proportion of the multiresistant ancestral strain RSDHI/DMI. The final concentration of *cytb^G143A^* measured in PopRR lines is indicated by a black crossed circle. Dashed gray lines represent the detection limit of the alleles. Interpretation of these figures and statistical models are detailed in **Supplementary Information 7** and **8**. ***A. Performance of strategies with poorly diverse selection pressure. B. Performance of strategies with highly diverse selection pressure*.**

By contrast, the introduction of a second AI with a different MoA (SDHI or DMI) significantly decreased population growth (by 2 - 85%, with a mean of 39%), whilst reducing the selection of resistant alleles (-49% - -100%, mean -82% for *sdhC^H152R^* and -43% - -100%, mean -87% for *cyp51^I381V^*, relative to selection with a single AI) in lines in which only single-resistant isolates to each of the two MoAs were initially introduced (**Fig. 6B** and **Supplementary Information 8**). The greatest reductions were observed with the P and B fungicides, rather than with the M and F ones. This benefit of combining MoAs was much more limited (from -4% to +4% of *M*^*n*^_*i*_, mean -0.6%) when the multiple-resistant isolate was introduced, due to its early and rapid generalization (increase by +85% - +596%; mean +241%), at the expense of single-resistant isolates, revealing its greater relative fitness when multiple selection is applied (**Fig. 4**, **Fig. 6B** and **Supplementary Information 5** and **8**). These findings confirm that diversifying modes of action, whatever the heterogeneity in the timing of their application (as a mixture or in alternation) is beneficial only in the absence of multiple resistance.

We investigated whether increasing the number of AIs and of MoAs induced the selection of new resistance profiles, by performing droplet tests on the 99 population lines evolved under one of the 31 selection regimes and collected at the end of the experiment, for 10 fungicide treatments and a control. These treatments included the fungicides used for selection but also other independent AIs (**Fig. 5C**; **Fig. 5D**; **Fig. 5E ; Supplementary Information 3**). Population resistance profiles were generally consistent with allele quantification, with the population displaying the resistance phenotype profile of the predominantly selected ancestral isolate. In particular, in PopRR lines that had evolved under diversified selection pressure from SDHIs combined with DMIs, low phenotypic diversity was observed, confirming the generalization of the multiple-resistant isolate. By contrast, greater diversity was observed after selection by a single MoA, reflecting the variation in resistance frequency of the single-resistant isolates between replicates and selection regimes. In PopR lines, the lack of growth in the presence of DMI and SDHI mixtures of lines evolved under a single selection pressure confirmed the preferential selection of single-resistant isolates. Interestingly, not only did lines evolved under diversified selection regimes grow in the presence of the two MoAs independently, confirming the co-selection of single-resistant isolates, but some also displayed additional resistances, as suggested by their greater diversity in the PCA and original patterns of resistance on heatmaps (**Fig. 5B**; **Fig. 5C**; **Fig.5D**). In particular, some lines selected under pressure from a combination of M (DMI) and F (SDHI) fungicides displayed resistance to the non-DMI and non-SDHI fungicides azoxystrobin and tolnaftate.

Tolnaftate has been shown to reveal multidrug resistance in *Z. tritici* (Leroux and Walker, 2011). We therefore checked for insertions in the promoter of the MFS1 transporter gene, which is overexpressed in field isolates (Omrane et al., 2017), in a selection of 20 strains isolated from these populations with various scores for tolnaftate resistance. We found no change in the length of the promoter sequence (data not shown). We then measured the expression of *mfs1* in the same isolates after growth in the absence of fungicide. We confirmed that nine of these isolates displayed moderately enhanced (about 3.3 - 10 times higher) fungicide efflux. By contrast, the level of expression of *mfs1* in the MDR field isolate was about 30 times higher, suggesting only moderately enhanced efflux due to MFS1 in some of our isolates, with still unknown determinism. Furthermore, the poor correlation between tolnaftate resistance scores and *mfs1* expression strongly suggests that unidentified generalist resistance mechanisms may be at work in our collection. Finally, this finding suggests that diversified selection pressure mediates a trade-off between the growth rate of the evolving population and the degree of generalism of the evolved isolates.

### The performance of strategies depends primarily on the diversity of MoAs displayed, but also on the particular characteristics of AIs

We further investigated the reasons for the better performance, in terms of effects on population growth, of strategies combining several MoAs and AIs than of continuous use of the same fungicide, by analyzing the effects of the respective intrinsic resistance risks of the MoAs and AIs on resistance evolution. This analysis was made possible by the use of two different MoAs (SDHIs *vs.* DMIs) and two AIs within each chemical class (B and F *vs.* P and M, respectively). It has been claimed that the intrinsic risk of resistance is low-to-moderate for DMIs, but moderate-to-high for SDHIs (Kuck and Russell, 2006). AIs were compared at the same effective dose (EC97) to ensure a fair comparison based on their intrinsic properties.

DMI-based strategies yielded a 4.4% higher *M*^*n*^_*i*_ mean normalized Malthusian growth than SDHI-based strategies, regardless of the initial population composition, the number of AIs in the strategy or the temporal heterogeneity of selection (**Fig. 6A** and **Supplementary Information 8**). This moderate difference suggests that the respective resistance risks of DMIs and SDHIs were not markedly different in the experimental conditions used here, at least for resistance dynamics, not anticipating field efficacy. In PopR, both MoAs were equally effective for reciprocally controlling the single-resistant isolates (the normalized concentration of *sdhC^H152R^* and of *cyp51^I381V^*after exposure to either DMIs or SDHIs, respectively, both below the detection limit) whereas in PopRR, the estimated normalized concentration of multiple resistance in the presence of DMIs was only half that with SDHIs only (16% *vs.* 33%), suggesting that SDHIs were less efficient than DMIs for controlling RSDHI/DMI. These findings mirror the contrasting selection responses, which are largely dependent on the resistance allele selected and its relative fitness. This finding is not, therefore, readily generalizable when diverse mutations may arise under selection pressure from a single MoA in the field, as is the case for SDHIs and DMIs.

The intrinsic resistance risk may also be AI-specific, and was therefore investigated for SDHIs and DMIs. In our conditions, the *M*^*n*^_*i*_ of strategies combining P and B was lower, by a factor of 1.45 - 5.7 (mean 2.4), than that of strategies combining M and F in PopR lines. This finding is consistent with the -55% to -100% decrease in the concentrations of *sdhC^H152R^* or *cyp51^I381V^* in strategies including P and B (**Fig. 6B** and **Supplementary Information 8**). The poorer control observed in M- and F-based strategies may also explain the emergence of additional generalist resistance mechanisms, as described above, in these lines (**Fig. 5D**). The benefit of strategies combining P and B over those combining M and F was also significant in PopRR lines, but to a much lesser extent (-2.6% of *M*^*n*^_*i*_; -1% in content of the multiple-resistant isolate). These findings suggest that it may also be difficult to generalize resistance risk to a whole MoA.

### New MoAs at appropriate doses are relevant partners for enhancing the performance of strategies

Fungicide diversity was found to affect the selection of resistance for AIs for which resistance was present at low frequencies in the initial populations. We also explored whether introducing a “new” AI, *i.e.* an AI to which the whole populations were fully susceptible, could improve the performance of strategies, and under which conditions. We therefore combined chlorothalonil (C), a multisite inhibitor with a very low risk of resistance, with the B and P fungicides previously used (**Fig. 6B** and **Supplementary Information 8**).

Introducing C into a combination of B and P in PopR lines induced a slight increase of *M*^*n*^_*i*_ (+46% for the BP/C alternation and +13% for the BPC mixture when compared with the already high-performing BP mixture), its overall value being close to the detection limit over the course of the experiment. This finding can be explained by equal full control of both single-resistant isolates by C that reinforces that of the BP mixture. By contrast, in PopRR, combining C with B and P had a beneficial effect on resistance (-48% and -30% *M*^*n*^_*i*_respectively, for the BP/C alternation and BPC mixture, compared with the poorly-performing BP mixture) and on the frequency of resistant alleles (-63% for *sdhC^H152R^*and -63% for *cyp51^I381V^* average). However, resistance dynamics differed depending on whether C was alternated or mixed with a BP mixture. In the mixture strategy, a half-dose of C was not sufficient to control the increasing proportion of the multiple-resistant isolate in the population and to compensate for the decrease in efficacy of P and B (*M*^*n*^_*i*_ = 0.66 and mean double resistant frequency = 38%). In the alternation strategy, C applied at full dose was fully effective against the RSDHI/DMI ancestral isolate, restored efficacy, and selected against multiple resistance every two cycles, enabling a better overall control of the population (*M*^*n*^_*i*_ = 0.49 and mean double resistant frequency = 25%). In populations in which resistance has emerged, the performance of strategies involving a new AI may therefore be dependent on its dose and its redundancy in the selection pattern.

In strategies combining B, C and P, the increase in MoA diversity due to the introduction of a third MoA could potentially increase the degree of generalism in the evolved populations, as previously described (Ballu et al., 2021; Ballu et al., 2023). This trade-off was not observed (*i.e.* no additional growth on tolnaftate; **Fig. 5E**) in PopRR lines where the multiple-resistant isolate was present and fitter, and invaded the population rapidly, at the expense of the susceptible isolate. However, PopR populations subjected to alternation or mixture involving C acquired generalist resistance, as revealed by their growth on tolnaftate, unlike all but one population subjected to a combination of B and P. This emergence and selection of generalist resistance might have been permitted by the maintenance of higher diversity in populations, more likely to experience favourable mutation. Therefore, when a new MoA is introduced and combined with previously used MoAs, attention must be paid to the prior presence or absence of certain resistances, which could strongly influence the risk of generalist resistance emerging, as well as specialist resistance to the new MoA, depending on its own resistance risk.

### Mixtures do not systematically outperform alternation: population composition also matters

As the search for the best anti-resistance strategies often pits mixtures against alternation, we studied the effect of temporal heterogeneity (alternating *vs.* mixing AIs) on resistance dynamics and phenotype evolution.

The temporal heterogeneity achieved by alternating instead of mixing AIs with the same MoA had no (for DMIs) or little (-3% to -5%; for SDHIs) effect on *M*^*n*^_*i*_, whatever the composition of the ancestral population (**Fig. 6A** and **Supplementary Information 8**). Within-MoA alternations and mixtures of SDHI and DMI fungicides led to the rapid generalization of *sdhC^H152R^* and *cyp51^I381V^*, respectively, in a similar manner. The rate of *sdhC^H152R^* selection was slightly lower (-9% in PopR and -2% in PopRR) for the alternation of B and F than for their mixture.

When SDHIs and DMIs were combined, their mixtures performed significantly but moderately better than their alternation (in average, -34% for *M*^*n*^_*i*_; -70% for *sdhC^H152R^* concentration; -99% for c*yp51^I381V^* concentration), only when single-resistant isolates were introduced (PopR lines). However, mixtures combining M and F were less effective than those combining B and P, because of the lower level of control exerted by their half-doses or differences in their inherent resistance risks, as explained above (**Fig. 6B** and **Supplementary Information 8**).

However, when the multiple-resistant isolate was present in the population (PopRR lines), it was rapidly and intensively selected by both alternation and mixture, resulting in poor performances for both types of strategy, for both *M*^*n*^_*i*_ and the selection of single-resistant isolates, although slight differences were observed between fungicide pairs. As described above, the addition of a third, new MoA (C) was more effective in alternations than in mixtures (-26% for *M*^*n*^_*i*_; -32% for maximal multiresistant isolate content) because it made it possible to use a full dose of C every second cycle.

Alternation and mixtures resulted in a diversity of resistance phenotype population profiles, reflecting the variable frequencies of the resistances to SDHIs or DMIs, or their co-occurrence (**Fig. 5D**; **Fig. 5E**). No resistances other than those initially introduced from the ancestral strains were observed, with the exception of the generalist resistance profiles of PopR lines, which were also able to grow on tolnaftate and azoxystrobin, as previously described. These generalist profiles were found exclusively in mixture and alternation selection regimes including two or three MoAs; neither alternation nor mixing could be considered more likely to favor generalist resistance. Temporal heterogeneity may not impact generalist resistance as much as the diversity of MoAs.

Overall, the advantage of mixtures over alternation selection regimes in terms of resistance dynamics and prevention of generalist resistance was driven mostly by the occurrence of multiple resistance in the ancestral population and the AIs used in the strategy and their doses.

## Discussion

This experimental study was designed to explore the performance of some of the drivers of resistance management strategies (*i.e.* the number and type of MoAs, the number and type of AIs, their vulnerability to resistance and the temporal pattern of fungicide exposure) relative to the resistance status of the population of the plant pathogenic fungus *Zymoseptoria tritici*. We ran a comprehensive experimental evolution study that was designed to disentangle the respective influences of these drivers on the evolution of resistance in three populations of different initial compositions (a sensitive population, and two others, each including 5% fungicide-resistant individuals carrying either only single target-site resistance alleles or single and multiple target-site resistance alleles). We used multicriteria assessment (population growth rate, resistance allele content and pattern of cross-resistance in evolved populations) to evaluate the performance of 31 selection regimes. We found that strategies combining several MoAs exerted sufficiently diversified selection pressure to mitigate the selection of resistance if the initial populations were fully susceptible or included only single-resistant isolates at low frequency. By contrast, the introduction of multiple-resistant isolates abolished the beneficial impact of the best strategy drivers, indicating that population composition is the principal factor to be considered when tailoring anti-resistance strategies.

### Alternation can be as sustainable as mixture and may then reconcile resistance management and socioenvironmental expectations

Using an original experimental evolution approach, we show here that maximizing the diversity of selection while smartly combining drivers can effectively delay the evolution of resistance, as previously reported in empirical studies comparing a limited number of field or modeled situations, based on assumptions simplifying the biological reality (REX_Consortium, 2013; van den Bosch et al., 2014a). In particular, our findings confirm that selection was weakest for the combination of several MoAs (*i.e.* when maximizing inter-MoA diversity), and stronger when AIs with the same MoA were combined (*i.e.* when relying only on intra-MoA diversity).

Indeed, the exclusive use of DMI or SDHI fungicides, with one or two AIs, rapidly led to the establishment of resistance, associated with a loss of full control. This situation mirrors findings previously obtained with the same approach concerning the emergence of resistance in a naïve population (Ballu et al., 2023). Several studies have documented the poor efficacy of sequences of fungicides with the same MoA for controlling target-site resistance in *Z. tritici,* in the field or in models (Mavroeidi and Shaw, 2006; Hobbelen et al., 2013; Dooley et al., 2016a; Heick et al., 2017b; Hagerty et al., 2021). Indeed, AIs with the same MoA generally display positive cross-resistance, as confirmed by the phenotyping of our ancestral isolates. However, resistance intensity may differ between AIs for the same resistance allele, or between alleles for the same AI, reflecting the variation in AI affinity for the mutated target protein (e.g. Rehfus et al., 2018), ranging from strong to weak or a lack of resistance to some specific AIs. AIs exceptionally not or less affected by cross-resistance may appear of interest from a practical standpoint and could be preferred to mixtures of AIs exhibiting full cross-resistance. The strength of selection of such intra-MoA mixtures may depend on the degree of cross-resistance for the specific AIs and genotypes concerned. However, this benefit is strongly limited by the high diversity of resistance mutations and phenotypes in field populations and by the later emergence of mutations determining full cross-resistance (Leroux and Walker, 2011; Huf et al., 2018; Rehfus et al., 2018; Garnault et al., 2021). Our results suggest that although combinations of fungicides sharing the same MoA involve potentially very diverse AIs, with the entire range of authorized SDHIs and DMIs generally used in agricultural areas, they do not provide sufficiently diverse selection pressure and globally favor the selection of resistance on a long-term perspective.

The use of multiple MoAs, rather than AIs, is another option to maximize the diversity of selection, but this can be achieved either in mixture, with all generations of the pathogen submitted to the same combination of selection pressures, or in alternation, in which a few generations are subjected to a single selection pressure different from that imposed on their ancestors or progeny. The efficacy of mixtures is based on redundant killing and the low probability of *de novo* multiple resistance developing, but also possibly on higher dose rates (see below) (Ballu et al., 2021), whereas alternation may delay the build-up of resistance by spreading selection pressure out over longer periods, and by decreasing the rate of evolution of resistance to the fungicides alternated (Ballu et al., 2023). Many studies have compared the efficacy of these strategies for limiting the emergence of resistance and selection in fungi and other taxa (see reviews in REX_Consortium, 2013; Kim et al., 2014; van den Bosch et al., 2014a). Most, but not all, of these studies ranked mixtures above alternation in terms of performance in theoretical and empirical studies and simplicity of implementation, notably due to the availability of commercial ready-to-use mixtures. In most studies, mixtures were often defined as the simultaneous application of two MoAs, both used at their effective dose (van den Bosch et al., 2014a). In the field, this would lead to the quantities of fungicide used almost doubling. In this study, we propose an original and fair comparison of alternation and mixtures at constant environmental impact, *i.e.* with equal cumulative amounts of fungicide. Mixtures were, thus, defined as half the selection dose of each AI calculated for susceptible isolates. In this context, we found one situation where mixture surpassed alternation, two situations where both strategies where equivalent, and one situation where alternation overcame mixture. More particularly, the mixture and alternation of SDHIs and DMIs were equally effective for delaying the selection of resistance and controlling the growth of initially naïve populations. Mixtures moderately outperformed alternation when single-resistant isolates were present in the initial population, because the partial and possibly synergistic efficacies of both partners remained sufficient to control emerging single-resistant isolates and keep them at low frequency in the population. By contrast, in alternation, each single-resistant isolate alternatively escaped control when subjected only to a full dose of the corresponding AI every other cycle, because this strategy allowed to increase frequency of both isolates, alternatively. Performance was also determined by the AIs combined (*e.g.* at the chosen selection doses, M and F performed less well than P and B, whatever the strategy used). Mixtures and alternation were equally poor when the multiple-resistant isolate was present in the initial population and selected, as both AIs were ineffective, which rapidly established itself within the population. Interestingly, in this context, the introduction of a new MoA to which the whole population was susceptible was more valuable in alternation than in mixtures, as alternation allowed maximal efficacy against multiple resistance due to the application of a full dose every other cycle. In situations in which multiple resistance is observed, the benefits of alternation may depend more on the dose, frequency of use and vulnerability to resistance of the new agent introduced, than on the AIs alternated. Here, the new fungicide introduced was a multisite inhibitor with a very low resistance risk. The introduction of a new, previously unused fungicide may be considered with caution for a single-site fungicide with a greater inherent risk of resistance, the efficacy of which may decrease over time. Overall, our findings demonstrate that at similar and constrained total dose, mixtures may not always outperform alternation and are in line with a recent modelling study (Hardy, 2022) that concludes that the relative efficacies of these strategies clearly depend on the composition of the target population. In this sense, although the situations explored here are quite common in the field, it would be interesting to explore the robustness of these conclusions for other scenarios of population composition (e.g. in a fully susceptible population or with pre-existing resistance to one fungicide but not yet to the other, as this situation can be observed in field conditions). These findings might be also highly relevant in a context where the reduction of pesticide use, more achievable by using alternation than by using many current mixtures, constitutes a strong socioenvironmental expectation in an increasing number of countries. They finally represent an exciting perspective for future field investigations, where the unpredictable nature of the wider environment may pose greater challenges and exaggerate some to the inherent differences that may exist between the different MoAs used.

### The performance of strategies creating highly diversified selection pressure is limited by multiple and generalist resistance

Our analysis of the relevance of drivers of resistance management strategies for delaying the development of resistance unequivocally identifies population composition with regard to resistance as the primary determinant of the performance of strategies, ahead of the diversity of the MoAs applied and temporal heterogeneity. Multiple resistance, in particular, is an aggravating factor, as it is very poorly controlled by the fungicides that contributed to its selection, whether they were used alone or in combination. In strategies combining multiple MoAs, multiple-resistant isolates are preferentially selected over single-resistant ones, because they confer greater relative fitness under multidimensionnal selection (Corkley et al., 2022). This study confirms *in vivo* the results of modeling studies considering multiple resistance (Hobbelen et al., 2013; Tepekule et al., 2017). Finally, this experimental evolution approach also provides evidence that increasing the dimensionality of selection may favor generalist phenotypes over specialist ones, a question rarely explored so far, except for a recent experimental work (White et al., 2022) whose results tend to agree with ours, and that would deserve complementary theoretical investigation (Mérot, 2022).

Generalist fungicide resistance is also observed when non target-site resistance is selected, that is, when resistance mechanisms other than those encoded by mutations in the target-encoding genes are selected and confer wide patterns of cross-resistance to unrelated fungicides (Berman and Krysan, 2020; Hu and Chen, 2021). In an experimental design in which resistance is already present, we expected to see the direct selection of the resistance alleles, but at contrasting intensities, depending on the selection regime applied. Surprisingly, the analysis of population phenotypes at the end of the evolution experiment revealed the occurrence of *de novo* phenotypes in some lines. In populations into which only single-resistant isolates were present, additional resistance to azoxystrobin, a QoI not used for selection, and tolnaftate, an inhibitor of sterol-squalene epoxidase generally used to reveal enhanced fungicide efflux (Leroux and Walker, 2011), formerly absent, was detected, exclusively in some lines subjected to alternations or mixtures of SDHIs and DMIs, associated or not to the multisite inhibitor C. This generalist resistance co-existed, within populations, with the initially introduced DMI and SDHI resistances. Generalist resistance can even be associated with target-site resistance allele, as highly probable in lines undergoing M and F selection where *sdhC^H152R^* was fully established at the end of the experiment. The purification of isolates resistant to tolnaftate made it possible to confirm the slight-to-moderate overexpression of *mfs1*, encoding a MFS transporter responsible for multidrug resistance in *Z. tritici* in the field (Omrane et al., 2017), in some, but not all of the strains tested. The absence of the usual regulatory insertions in the promoter of *mfs1*, and the poor correlation between susceptibility to tolnaftate and *mfs1* expression suggest that unknown mutations and resistance mechanisms probably underlie the generalist resistance of these isolates. This evolutionary trade-off in resistance management, with attempts to reduce selection for specialist resistance traits promoting the evolution of generalist resistance, was observed in a previous study dealing with alternation and effective-dose mixture strategies in naïve populations (Ballu et al., 2021; Ballu et al., 2023). It may apply more generally to any strategy combining multiple MoAs in alternation or mixtures and aiming to mitigate the evolution of resistance in other organisms, such as weeds (Lagator et al., 2013a; Lagator et al., 2013b; Comont et al., 2020) or bacteria (Kim et al., 2014). This trade-off is still little considered in the field, with attention instead focused on more quantitative criteria for measuring strategy performance, such as resistance frequency, or the effective life-time of a fungicide. However, high frequencies of individuals with generalist resistance in the pathogen population may irreversibly decrease the number of effective AIs even more than TSR. In weeds and insects, NTSR and TSR generally occur together and are associated with high RFs and unexpected resistance patterns (Mitchell et al., 2014; Gaines et al., 2020). In *Z. tritici,* MDR *per se* is associated with moderate resistance factors, but its combination with target-site alterations has been found to increase resistance factors to DMIs or SDHIs significantly (unpublished data).

Given the importance of population composition for strategy performance and the trade-off described above, these findings suggest that the debate should be refocused on the prevention of multiple and, to a lesser extent, generalist resistance, rather than on the relative merits of mixtures and alternation, as summarized in (Corkley et al., 2022).

### The crux of resistance management should focus on limiting multiple resistance

Multiple resistance occurs in populations for several reasons. First, it reflects the sequential use of MoAs, following their successive discoveries and withdrawals following the development of intensive cropping systems, which began just after World War II. Resistance led to new AIs often being combined with their predecessors to improve residual efficacy, and resistance to new MoAs was successively selected in genetic backgrounds that were already resistant to the older MoAs, which were generally withdrawn once resistance became widely established. Multiple resistance has been described as resulting from a stepwise accumulation of resistances in several plant pathogens (*e.g. Venturia inaequalis* (Koller and Wilcox, 2001), *Monilinia fructicola* (Luo and Schnabel, 2008), *Botrytis cinerea* (Li et al., 2014), and *Cercospora beticola* (Secor et al., 2010)), and it may even favor the development of resistance to new MoAs. *Z. tritici* is no exception, as resistance to benzimidazoles and QoIs remains widespread in populations from Western Europe, despite the withdrawal of these fungicides from the market to control septoria leaf blotch (although QoIs are still little used against other diseases). The ongoing selection of resistance to SDHIs is almost systematically associated with

DMI resistance, simply because it is selected within DMI-resistant populations (Dooley et al., 2016b; Garnault et al., 2019).

Second, multiple resistance may result from the recombination of resistance alleles during sexual reproduction. The primary inoculum of *Z. tritici,* including ascospores, is never a limiting factor for epidemics and sexual reproduction. Indeed, population-level deep sequencing revealed intense interplay between asexual and sexual reproduction (Singh et al., 2021). In this airborne pathogen, the ascospores resulting from sexual reproduction are dispersed over long distances and have been reported to spread QoI resistance between fields locally (Fraaije et al., 2005) but also at the continental scale in Europe (Torriani et al., 2009).

Despite these strong assumptions on the occurrence of multiple resistance in agricultural systems, the performance of strategies has generally, with few exceptions, been modeled from initially susceptible or populations containing single-resistant isolates. Yet, the reality in the field is that new AIs are probably only rarely confronted with the ideal situation in which all the AIs and MoAs used are fully effective. In the real world, multiple resistance is the norm, and the initial frequency of single- and multiple-resistant isolates almost certainly differ at fine scales depending on the historical use of fungicides and resistance management ; strategies need to be adapted accordingly (Taylor and Cunniffe, 2022). In this work, we assessed the performance of strategies in conditions mimicking real-life conditions, while artificially introducing single- and multiple-resistant isolates, to reproduce the combination of resistance alleles likely to occur in the field by sexual reproduction, but hardly achievable in the laboratory. However, the lack of this phenomenon in our experimental system precludes any assessment of the time required for the emergence of multiple resistance by recombination, all the more since preexisting resistances are often established by stepwise accumulation in populations.

One of the first conclusions to be drawn from our findings is that the design of resistance management strategies, and the long-term durability of MoAs would greatly benefit from resistance monitoring data (*i.e.* the composition of the local population) being made publicly available at the finest possible scale and detection limit. The high-throughput quantification of multiple resistance genotypes may become possible in the near future with the development of next-generation sequencing techniques for use in diagnostics for plant pathogen populations (Radhakrishnan et al., 2019; Samils et al., 2021). This, in turn, should make it possible to take informed decisions in spraying programs. Second, more attention should be paid to limiting the development of multiple resistance, and of the combination of TSR and NTSR, as both could greatly decrease the efficacy of any strategy. Recombination in airborne pathogens such as *Z. tritici* is an aggravating factor, but it is one that is not really preventable. However, delaying the emergence of single-resistant isolates and limiting the probability of recombination between them by ensuring that their frequencies remain low might be a more realistic option. It has been estimated that *Z. tritic*i produces at least 2.1-9.9 trillion pycnidiospores per hectare (McDonald et al., 2022). The number of *de novo* mutation events associated with resistance occurring in a field may, therefore, be sufficiently high to prevent all of these mutations being lost by chance, even if the mutation rate is low. The time to the emergence of full resistance has been found to decrease monotonically with increasing dose, over realistic ranges of pathogen life-history and fungicide dose-response parameters (Mikaberidze et al., 2017). The maintenance of low frequencies of resistance alleles during the selection phase would involve limitations on the use of all MoAs, especially when the frequency of resistance reaches a critical threshold over a large area. Disease control should more than ever be based on complementarity with other control measures, such as the replacement of synthetic fungicides by biological agents, resistant cultivars, prophylactic measures and agricultural practices, to increase the diversity of selection pressures and limit the emergence of resistance by controlling population size. The overall sustainability of integrated pest management may therefore rely on the vulnerability of each method to fungal adaptation and then on our capacity to predict and monitor these phenomena. This would also lead the community to consider criteria other than the effective life of fungicides when assessing the durability of crop protection. Assessing qualitative changes in evolved population while evaluating the diversity of protection measures used would also help us to work towards current socioenvironmental expectations while better reflecting the multiple evolutionary processes at work. However, although adapting the way we conceive resistance management strategies to focus more on multiple resistance would be a huge step forward in itself, the implementation of these strategies in practice will probably prove an even greater challenge. Indeed, optimal disease control would imply using the best fungicides and at high rates in standardized programs. By contrast, resistance management with limited environmental impact would imply using the whole range of available AIs, whatever their intrinsic efficacy, within parsimonious combinations tailored to the current local composition of the population, associated with other control measures. These approaches are conceived for short (*i.e.* an agricultural season) and long (*i.e.* decades) time scales, respectively, and maximize profit or promote sustainable agriculture, respectively, potentially creating an intractable dilemma for growers. Education about resistance issues and communication between stakeholders and farmers are almost certainly the key to changing the paradigm and initiating a transition towards agroecology.

## Data availability

Data and scripts were deposited on the Research Data Gouv repository under DOI https://doi.org/10.15454/DCVDZN.

Citation : Ballu, A., et al. (2022). Dataset of an experimental evolution on the effect of fungicide-resistance composition of populations on the durability of anti-resistance strategies on the wheat pathogen *Zymoseptoria tritici*, Recherche Data Gouv, https://doi.org/10.15454/DCVDZN.

## Supporting Information Captions

Supplementary information 1: Phenotype of ancestral isolates

Supplementary information 2: Selection regimes used for experimental evolution

Supplementary information 3: Doses of fungicides used for experimental evolution Supplementary information 4: Molecular quantification of resistance alleles in evolved lines

Supplementary information 5: cytb^G143A^ content of evolved lines for PopRR

Supplementary information 6: Expression of *mfs1* in evolved lines of *Zymoseptoria tritici*

Supplementary information 7: Models used for data analysis

Supplementary information 8: Performance of management strategies according to the selection regime applied to the ancestral population of *Z. tritici*.

## Acknowledgments

We thank Fabrice Blanc for facilitating the administrative organization of this PhD studentship. We would also like to thank Dr. Gabriel Scalliet, Dr. Stephanie Bedhomme and Dr. Mato Lagator for their sound comments about our findings.

## Author contributions

AB, ASW, FC, AD and CU conceived and designed the study. AB and CU performed the experimental evolution experiment with the help of CD. CU, AB and CD isolated evolved strains and established their phenotypes and genotypes. AN analysed the expression of *mfs1* in evolved isolates. JW and ST quantified alleles content in evolved populations. AB and FC performed the statistical analysis, with contributions from ASW and AD. The paper was written by AB, ASW, FC and AD, with additional contributions from all authors.

## Supplementary information 1: Phenotype of ancestral isolates

**Supplementary Table 1:** Susceptibility of ancestral isolates to various fungicides. The susceptibility of the ancestral isolates is expressed as the EC50s (mg.L^-1^) calculated from dose-response curves established with cultures in microtiter plates and a logistic non-linear mixed-effects model. Dose-response curves were plotted one to five times, depending on the AI (number of replicates specified in brackets), with four technical replicates. The standard error is specified in brackets.

| $EC_{50}$ (mg.L<sup>-1</sup>) | S | R<sub>SDHI</sub> | R<sub>DMI</sub> | R<sub>SDHI</sub>/DMI |
| --- | --- | --- | --- | --- |
| Benzovindiflupyr | 0.028<br>(±0.003) (n=2) | 2.556<br>(±0.407) (n=3) | 0.103<br>(±0.008) (n=2) | 6.84<br>(±5.224) (n=4) |
| Boscalid | 0.242<br>(±0.013) (n=4) | >100<br>(n=1) | 0.37<br>(±0.025) (n=4) | >100<br>(n=1) |
| Fluxapyroxad | 0.053<br>(±0.005) (n=3) | >50<br>(n=1) | 0.152<br>(±0.024) (n=3) | >50<br>(n=1) |
| Mefentrifluconazole | 0.001<br>(±0) (n=3) | 0.001<br>(±0) (n=3) | 0.009<br>(±0.001) (n=2) | 0.013<br>(±0.003) (n=3) |
| Metconazole | 0.004<br>(±0.001) (n=2) | 0.004<br>(±0) (n=2) | 0.087<br>(±0.015) (n=2) | 0.132<br>(±0.003) (n=1) |
| Prothioconazole-desthio | 0.002<br>(±0) (n=3) | 0.001<br>(±0) (n=3) | 0.042<br>(±0.004) (n=3) | 0.045<br>(±0.009) (n=3) |
| Carbendazim | 0.115<br>(±0.037) (n=3) | >50<br>(n=3) | >50<br>(n=3) | >50<br>(n=3) |
| Azoxystrobin | 0.047<br>(±0.004) (n=4) | 0.029<br>(±0.004) (n=3) | 0.033<br>(±0.004) (n=5) | 21.401<br>(±37.046) (n=4) |
| Chlorothalonil | 1.185<br>(±0.104) (n=3) | 0.494<br>(±0.021) (n=2) | 0.789<br>(±0.061) (n=3) | 0.927<br>(±0.076) (n=2) |
| Tolnaftate | 0.23<br>(±0.022) (n=4) | 0.338<br>(±0.068) (n=3) | 0.307<br>(±0.036) (n=4) | 0.116<br>(±0.011) (n=3) |

## Supplementary information 2: Selection regimes used for experimental evolution

**Supplementary Table 2:** Selection regimes and their respective components and codes. Thirty-one selection regimes were applied to three artificial populations of *Z. tritici —* PopS, PopR and PopRR — as defined in **Table 1A**. These regimes were organized into two batches (A and B). The selection regimes differed in their components, which defined the diversity and heterogeneity of selection. Control regimes (not including fungicides) were amended with 0.5% ethanol or DMSO (*i.e.* the amount of fungicide solvent used in treated lines). Straight selection regimes involved the continuous use of a single fungicide. The regimes displaying diverse selection included two or three fungicides differing in their inherent resistance risk and mode of action, and their temporal heterogeneity over the course of the experiment.

| Population tested | Number of AIs |  | Number of MoAs | Vulnerability to resistance<br>(resistance risk) |  |  |  |  | Temporal pattern of fungicide application |  | Name of the selection regime | Experimental batch |
| --- | --- | --- | --- | --- | --- | --- | --- | --- | --- | --- | --- | --- |
|  | Number of fungicides | Includes a multisite inhibitor |  | Medium/High<br>(SDHI) |  | Medium/Low<br>(DMI) |  | Very low<br>(multisite) | Intra-cycle (mixture) | Inter-cycle (alternation) |  |  |
|  |  |  |  | Benzovindiflupyr (B) | Fluxapyroxad (F) | Prothioconazole-desthio<br>(P) | Mefentrifluconazole (M) | Chlorothalonil (C) |  |  |  |  |
| PopS | 0 |  | 0 |  |  |  |  |  |  |  | PopS_Ethanol | A |
| PopS | 2 |  | 2 | X |  | X |  |  | X |  | PopS_Mixt_PB | A |
| PopS | 2 |  | 2 | X |  | X |  |  |  | X | PopS_Alt_PB | A |
| PopR | 0 |  | 0 |  |  |  |  |  |  |  | PopR_Ethanol | A/B |
| PopR | 0 |  | 0 |  |  |  |  |  |  |  | PopR_DMSO | B |
| PopR | 1 |  | 1 | X |  |  |  |  |  |  | PopR_B | A |
| PopR | 1 |  | 1 |  |  | X |  |  |  |  | PopR_P | A |
| PopR | 2 |  | 1 | X | X |  |  |  | X |  | PopR_Mixt_BF | B |
| PopR | 2 |  | 1 | X | X |  |  |  |  | X | PopR_Alt_BF | B |
| PopR | 2 |  | 1 |  |  | X | X |  | X |  | PopR_Mixt_MP | B |
| PopR | 2 |  | 1 |  |  | X | X |  |  | X | PopR_Alt_MP | B |
| PopR | 2 |  | 2 | X |  | X |  |  | X |  | PopR_Mixt_PB | A |
| PopR | 2 |  | 2 | X |  | X |  |  |  | X | PopR_Alt_PB | A |
| PopR | 2 |  | 2 |  | X |  | X |  | X |  | PopR_Mixt_MF | B |
| PopR | 2 |  | 2 |  | X |  | X |  |  | X | PopR_Alt_MF | B |
| PopR | 3 | X | 3 | X |  | X |  | X | X |  | PopR_Mixt_CPB | B |
| PopR | 3 | X | 3 | X |  | X |  | X | X | X | PopR_Alt_CPB | B |
| PopRR | 0 |  | 0 |  |  |  |  |  |  |  | PopRR_Ethanol | A/B |
| PopRR | 0 |  | 0 |  |  |  |  |  |  |  | PopRR_DMSO | B |
| PopRR | 1 |  | 1 | X |  |  |  |  |  |  | PopRR_B | A |
| PopRR | 1 |  | 1 |  |  | X |  |  |  |  | PopRR_P | A |
| PopRR | 2 |  | 1 | X | X |  |  |  | X |  | PopRR_Mixt_BF | B |
| PopRR | 2 |  | 1 | X | X |  |  |  |  | X | PopRR_Alt_BF | B |
| PopRR | 2 |  | 1 |  |  | X | X |  | X |  | PopRR_Mixt_MP | B |
| PopRR | 2 |  | 1 |  |  | X | X |  |  | X | PopRR_Alt_MP | B |
| PopRR | 2 |  | 2 | X |  | X |  |  | X |  | PopRR_Mixt_PB | A |
| PopRR | 2 |  | 2 | X |  | X |  |  |  | X | PopRR_Alt_PB | A |
| PopRR | 2 |  | 2 |  | X |  | X |  | X |  | PopRR_Mixt_MF | B |
| PopRR | 2 |  | 2 |  | X |  | X |  |  | X | PopRR_Alt_MF | B |
| PopRR | 3 | X | 2 | X |  | X |  | X | X |  | PopRR_Mixt_CPB | B |
| PopRR | 3 | X | 2 | X |  | X |  | X | X | X | PopRR_Alt_CPB | B |

## Supplementary information 3: Doses of fungicides used for experimental evolution

**Supplementary Table 3A:**
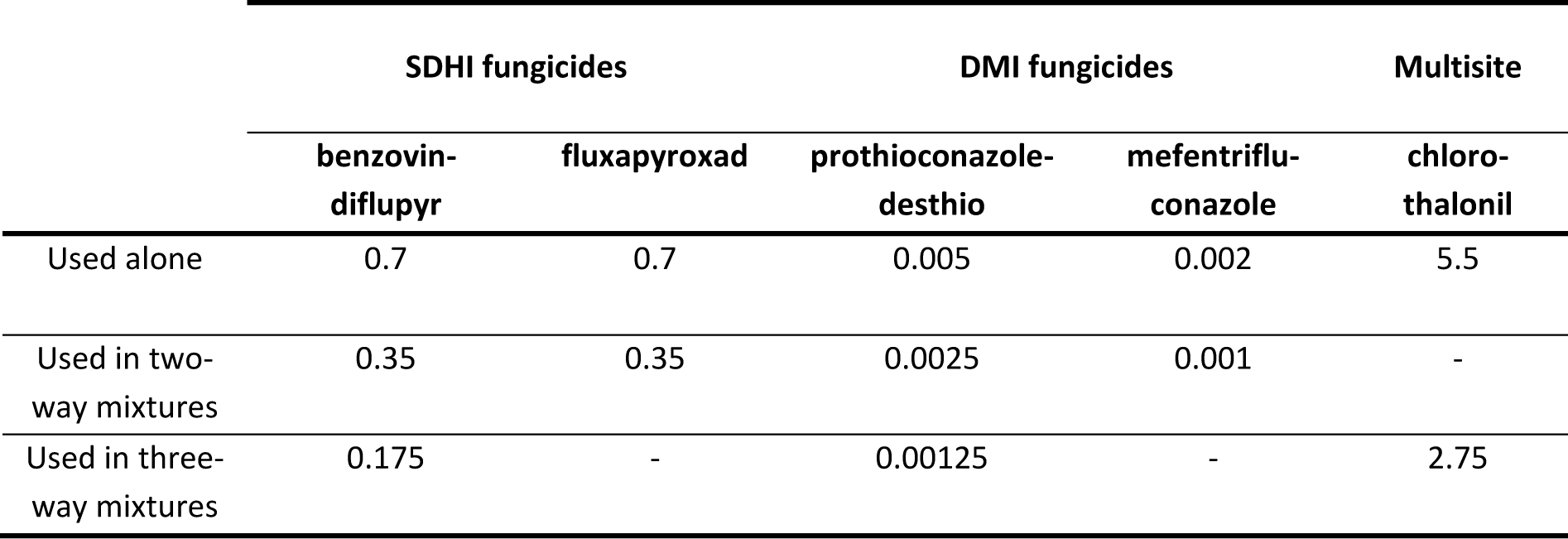
Selection doses of fungicides used for experimental evolution (mg.L^-1^).

|  | SDHI fungicides |  | DMI fungicides |  | Multisite |
| --- | --- | --- | --- | --- | --- |
|  | benzovin-<br>diflupyr | fluxapyroxad | prothioconazole-desthio | mefentriflu-<br>conazole | chloro-<br>thalonil |
| Used alone | 0.7 | 0.7 | 0.005 | 0.002 | 5.5 |
| Used in two-way mixtures | 0.35 | 0.35 | 0.0025 | 0.001 | - |
| Used in three-way mixtures | 0.175 | - | 0.00125 | - | 2.75 |

**Supplementary Table 3B:** Discriminatory doses used in droplet tests (mg.L^-1^).

| Acronym in Fig. 5 | Fungicides | Fungicide<br>class | Dose |
| --- | --- | --- | --- |
| B | benzovindiflupyr | SDHI | 0.7 |
| F | fluxapyroxad | SDHI | 0.7 |
| FB | fluxapyroxad + benzovindiflupyr | SDHI+SDHI | 0.7+0.7 |
| P | prothioconazole-desthio | DMI | 0.005 |
| M | mefentrifluconazole | DMI | 0.002 |
| MP | mefentrifluconazole + prothioconazole-desthio | DMI+DMI | 0.002+0.005 |
| PB | prothioconazole-desthio + benzovindiflupyr | DMI+ SDHI | 0.005+0.7 |
| MF | mefentrifluconazole + fluxapyroxad | DMI+ SDHI | 0.002+0.7 |
| Azo | azoxystrobin | QoI | 0.5 |
| Tol | tolnaftate | SBI class IV | 1.5 |
| C | chlorothalonil | Multisite | 5.5 |

## Supplementary information 4: Molecular quantification of resistance alleles in evolved lines

DNA was extracted from samples stored in microtiter plates with the DNeasy® Plant Kit (Qiagen), according to the manufacturer’s procedure.

The *sdhC^H152R^* allele, included in the RSDHI and RSDHI/DMI ancestral isolates, was quantified by qPCR. Allele-specific primers were designed with the goal of amplifying only this allele, with a primer discriminating between this and all other known alleles. The allele-specific forward primer CGT TGA ATG GAG TGA GGC A and the reverse primer TGT ACC ATC TCT CTT CAT CCT C amplified the product of the wild-type allele H152, whereas the allele-specific forward primer GTT GAA TGG AGT GAG GCG, used with the same reverse primer, amplified the product of the mutant allele R152. The non-specific forward primer TCG TTG AAT GGA GTG AGG C was used to estimate the concentrations of the reference gene (wildtype, mutant). The reactions were performed with FastStart Universal SYBR Green Master (ROX) (Roche Diagnostics GmbH, Roche Applied Science, Mannheim, Germany) on a Bio-Rad CFX-384 machine. The PCR was run under the following conditions: initial denaturation at 95°C for 10 min, 40 cycles at 95°C for 15 s, 58°C for 30 s and 75°C for 30 s (measurement of fluorescence). A melting curve, from 65°C to 95°C was recorded after the PCR. The detection threshold was estimated at 1%.

The *cyp51^I381V^*allele, included in the RDMI and RSDHI/DMI ancestral isolates, was quantified by pyrosequencing. A 216 bp fragment containing the 381 codon was amplified with the primer pair CCC GAC ATC CAA GAC GAA C and Biotin-TGG AAT GAC GTA TGC CGT ACC. The PCR conditions were as follows: initial denaturation at 95°C for 2 min, 50 cycles of 95°C for 30 s, 58°C for 30 s and 72°C for 30 s and a final elongation at 72°C for 5 min (GoTaq® Hot Start Polymerase, Promega, Madison, WI). We prepared 20 µL of PCR product for pyrosequencing reactions with the PyroMark Q96 Vacuum Workstation (Qiagen) and Streptavidin Sepharose High-Performance Beads (GE Healthcare Bio- Sciences AB, Uppsala, Sweden), as described in the workstation instructions and in the PyroMark Q96 ID user manual. The single-stranded DNA templates were transferred to 40 µL annealing buffer (Qiagen) containing the sequencing primer. The primer used in the reaction was ACC CTT CGT ATT CAC G [0.4 µM] for sequencing the codon 381 mutation site. The sequencing run was set up with PyroMark- Software v.1.0 (Qiagen). Samples were incubated at 80° C for 2 min and then allowed to cool to room temperature. The sequencing reaction was then performed with PyroMark Gold Q96 reagents on a PyroMark Q96 ID machine (both from Qiagen). The dispensation order of the nucleotides was AGC GTC AGA TCA. The detection threshold was estimated at 5% for the I381V substitution in preliminary experiments.

The *cytb^G143A^* allele, included only in the RSDHI/DMI ancestral isolate was quantified by qPCR. The forward primers were allele-specific discriminating for the wild type allele G143 (ACC TTA TGG TCA AAT GTC TTT ATG ATG) and for the mutant allele A143 (ACC TTA TGG TCA AAT GTC TTT ATG ATC). The conserved reverse primer used was AGC AAA GAA TCT GTT CAA TGT TGC. The reactions were performed with FastStart Universal SYBR Green Master (ROX) (Roche Diagnostics GmbH, Roche Applied Science, Mannheim, Germany) on a Bio-Rad CFX-384 machine. The PCR was run under the following conditions: initial denaturation at 95°C for 10 min, 40 cycles at 95°C for 15 s, 60°C for 30 s and 71°C for 30 s (measurement of fluorescence). The detection threshold was estimated at 1%.

## Supplementary information 5: *cytb^G143A^* content of evolved lines for PopRR

**Supplementary Figure 5:**
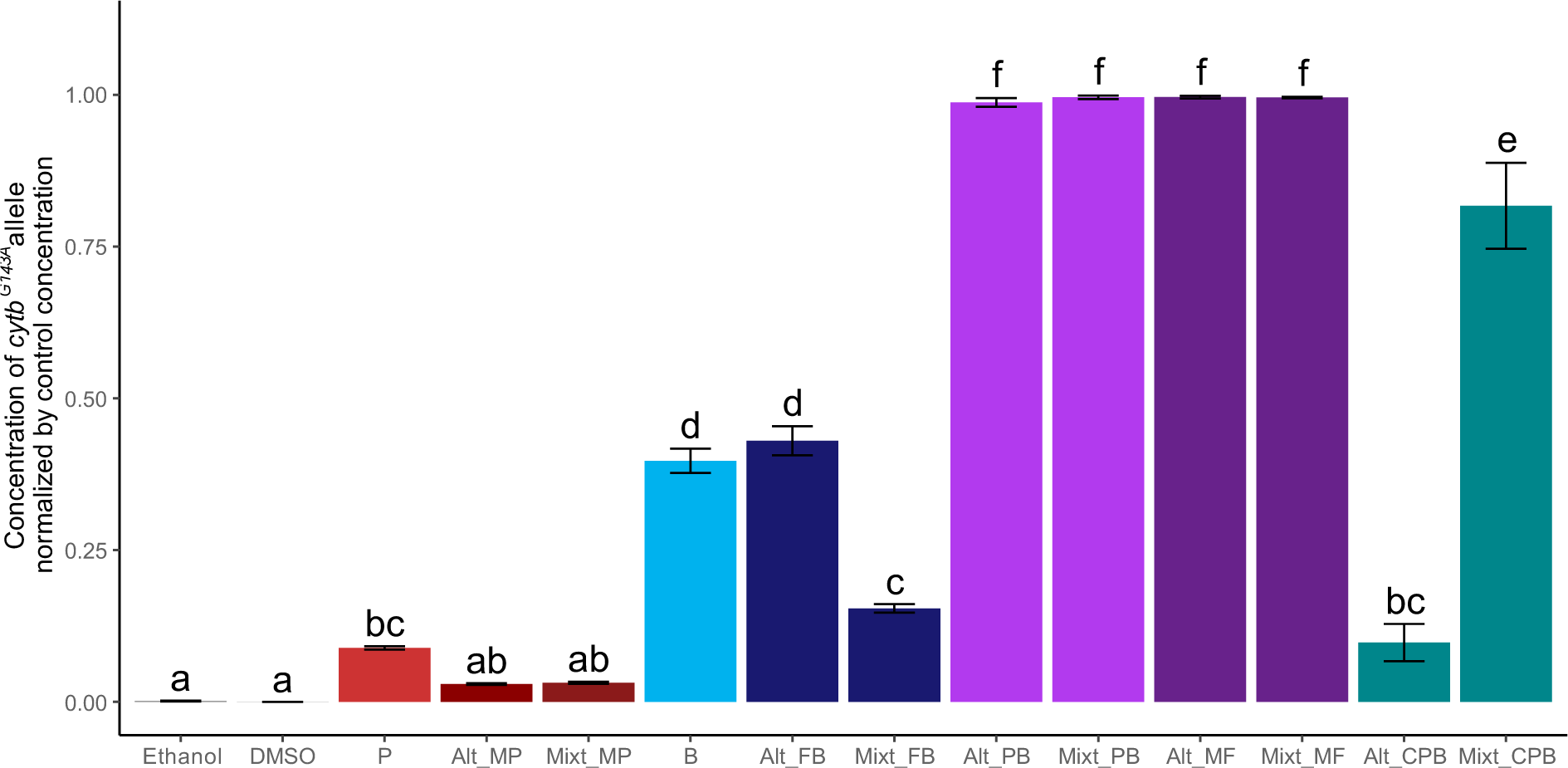
Normalized concentration of *cytb^G143A^* in lines evolved from the PopRR ancestral population. *cytb^G143A^* confers high-level resistance to QoI fungicides (not used in the strategies compared here). It was quantified by qPCR when optical density reached 90% of that of the control for a minimum of three consecutive cycles (4^th^, 6^th^ or 8^th^ cycle, depending on the line). This allele was carried exclusively by the RSDHI/DMI ancestral isolate and was used for indirect quantification of the proportion of isolates displaying multiple resistance in PopRR lines. Indeed, Pearson’s correlation coefficient for the relationship between the concentration of *cytb^G143A^*and the estimated concentration of multiple-resistant isolate (Fig. 4) in the same population was 0.998 (*P*<0.0001). Pairwise comparisons of lines evolved under different selection regimes were adjusted with the Tukey method for multiple comparisons (MODG143A0; SI2). The multiple-resistant was preferentially selected in lines subjected to diversified SDHI and DMI selection as opposed to straight selection under a single fungicide or the mixture or alternation of fungicides with the same mode of action (P<0.001; MODG143A1). There was, therefore, a resistance cost associated with the RSDHI/DMI ancestral isolate in competition with isolates carrying single resistance alleles in the presence of homogeneous selection pressure.

## Supplementary information 6: Expression of *mfs1* in evolved lines of Zymoseptoria tritici

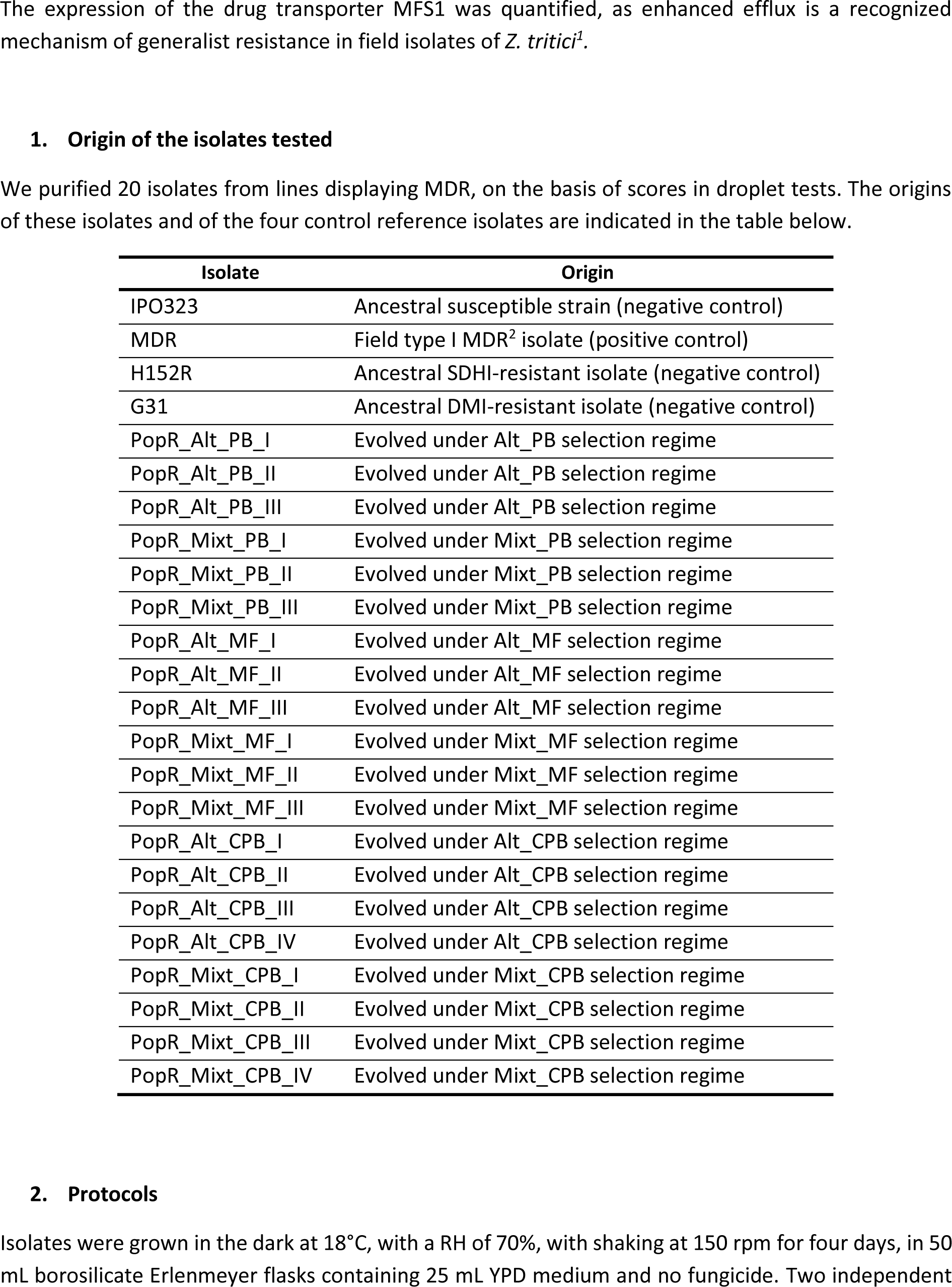
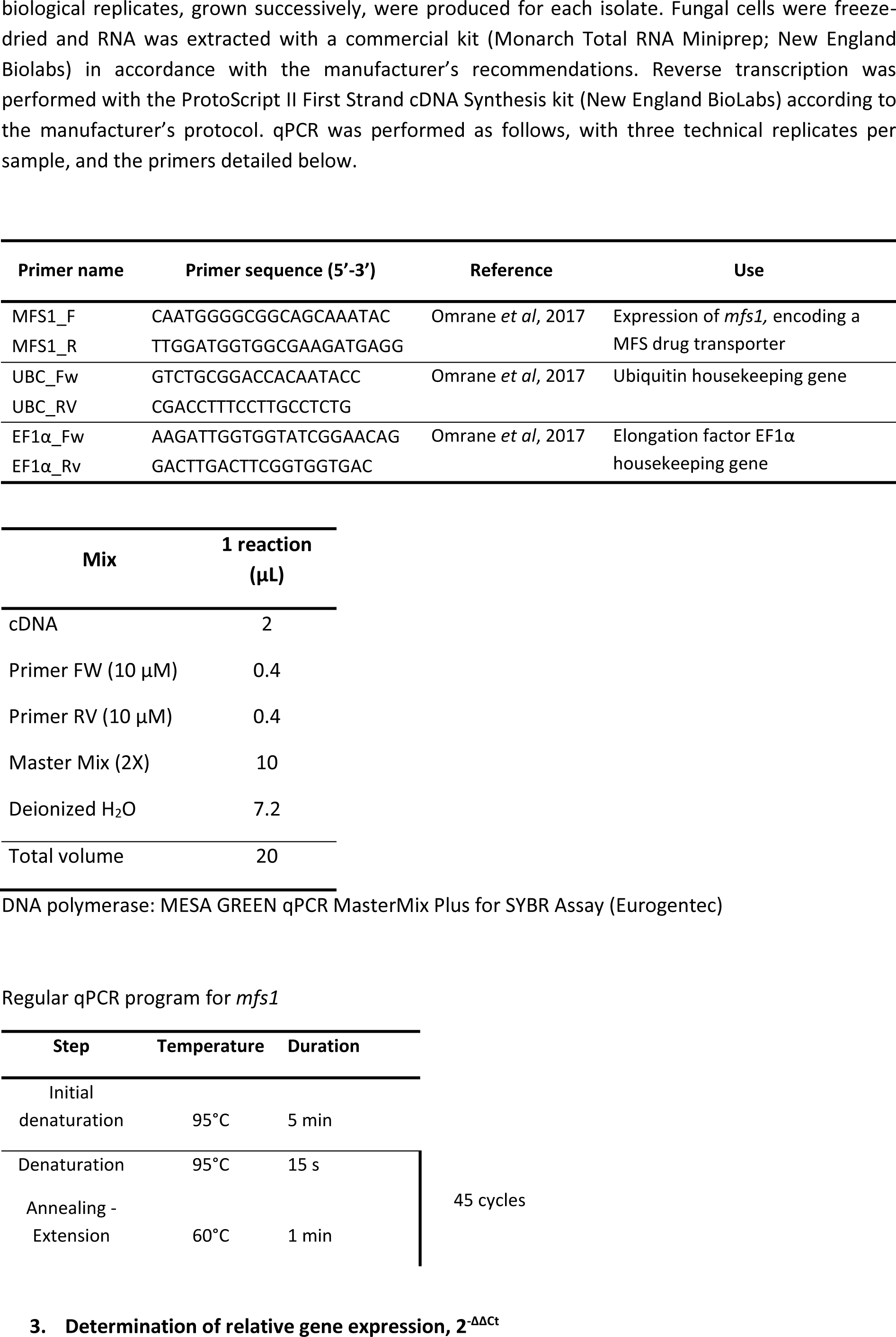
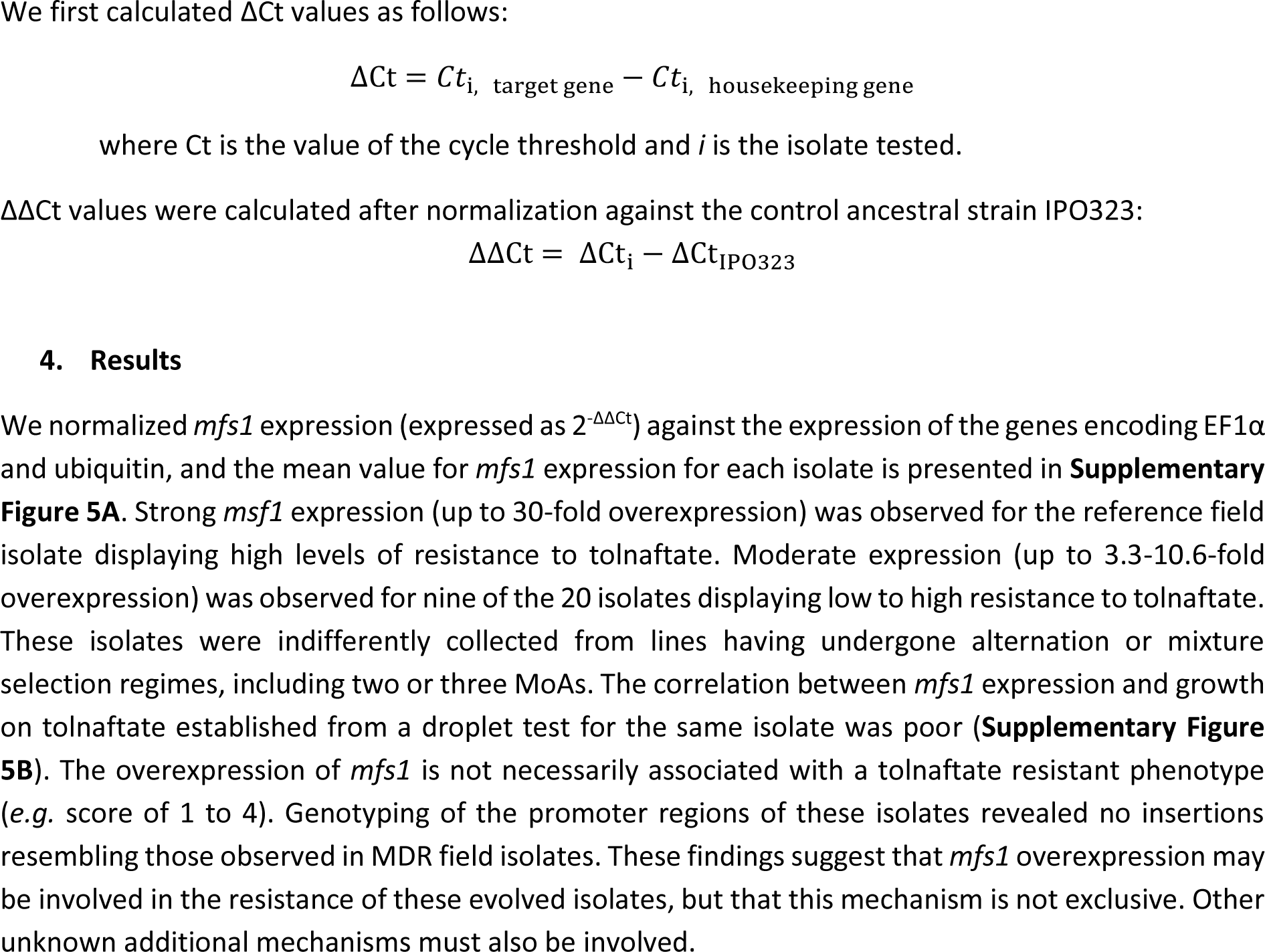
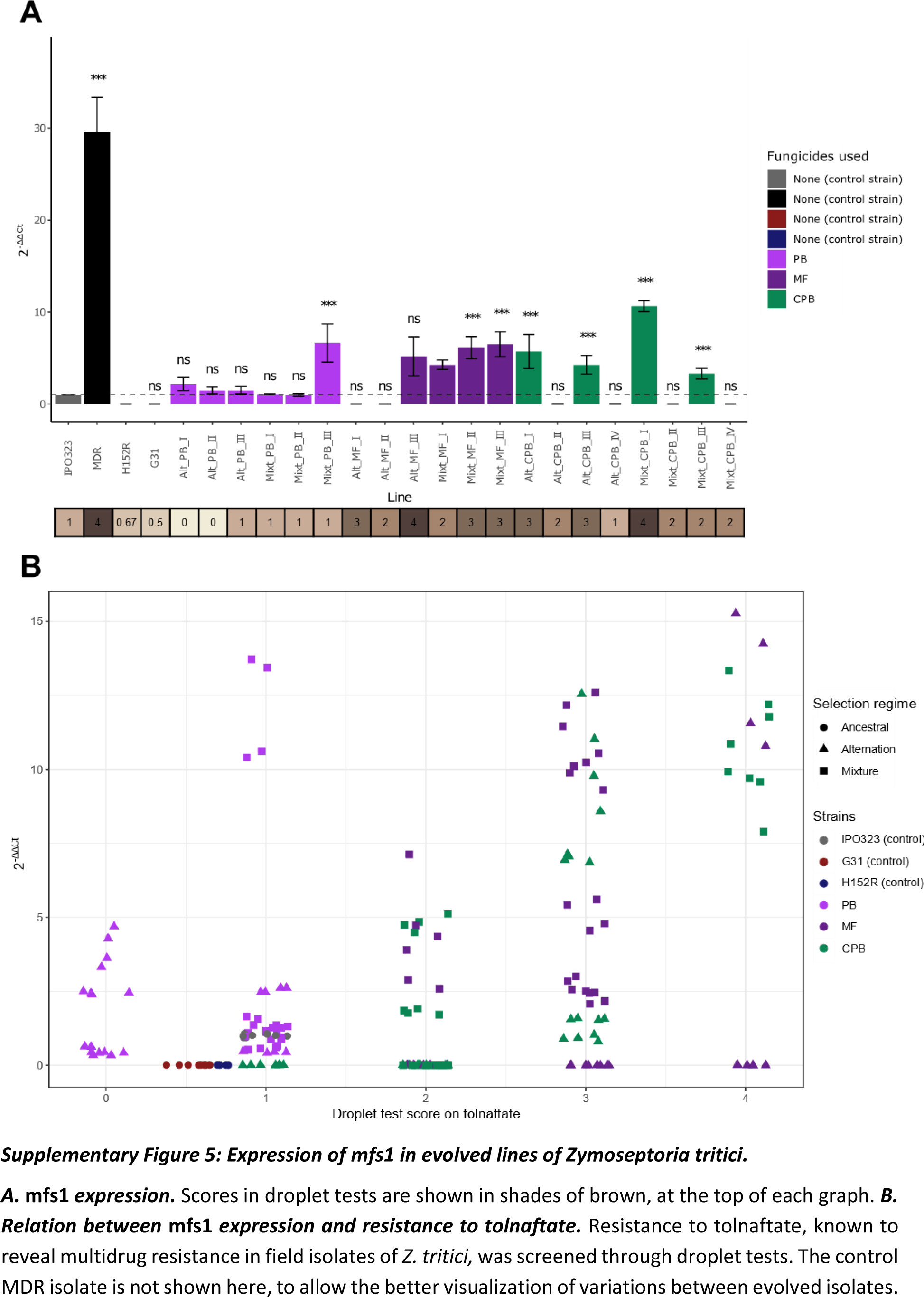
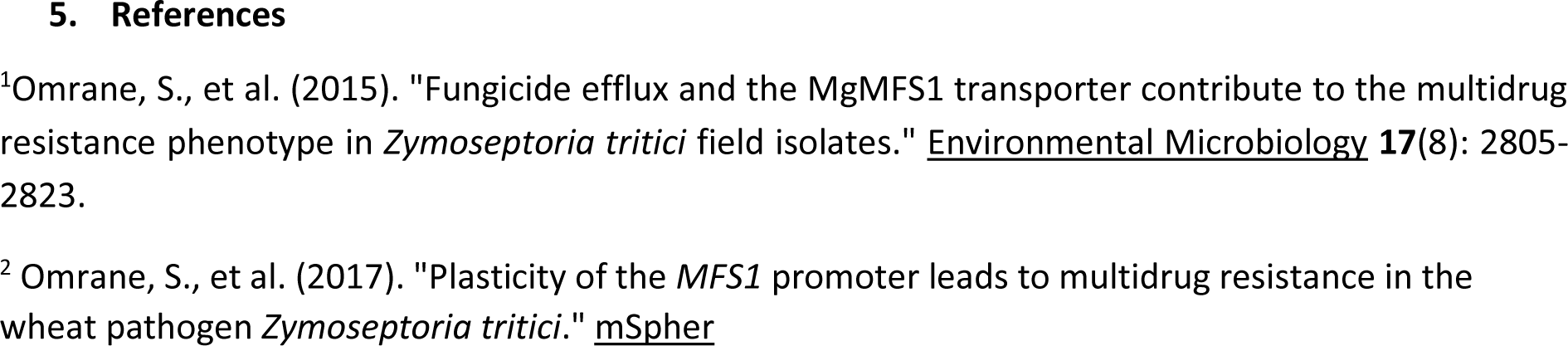

## Supplementary information 7: Models used for data analysis

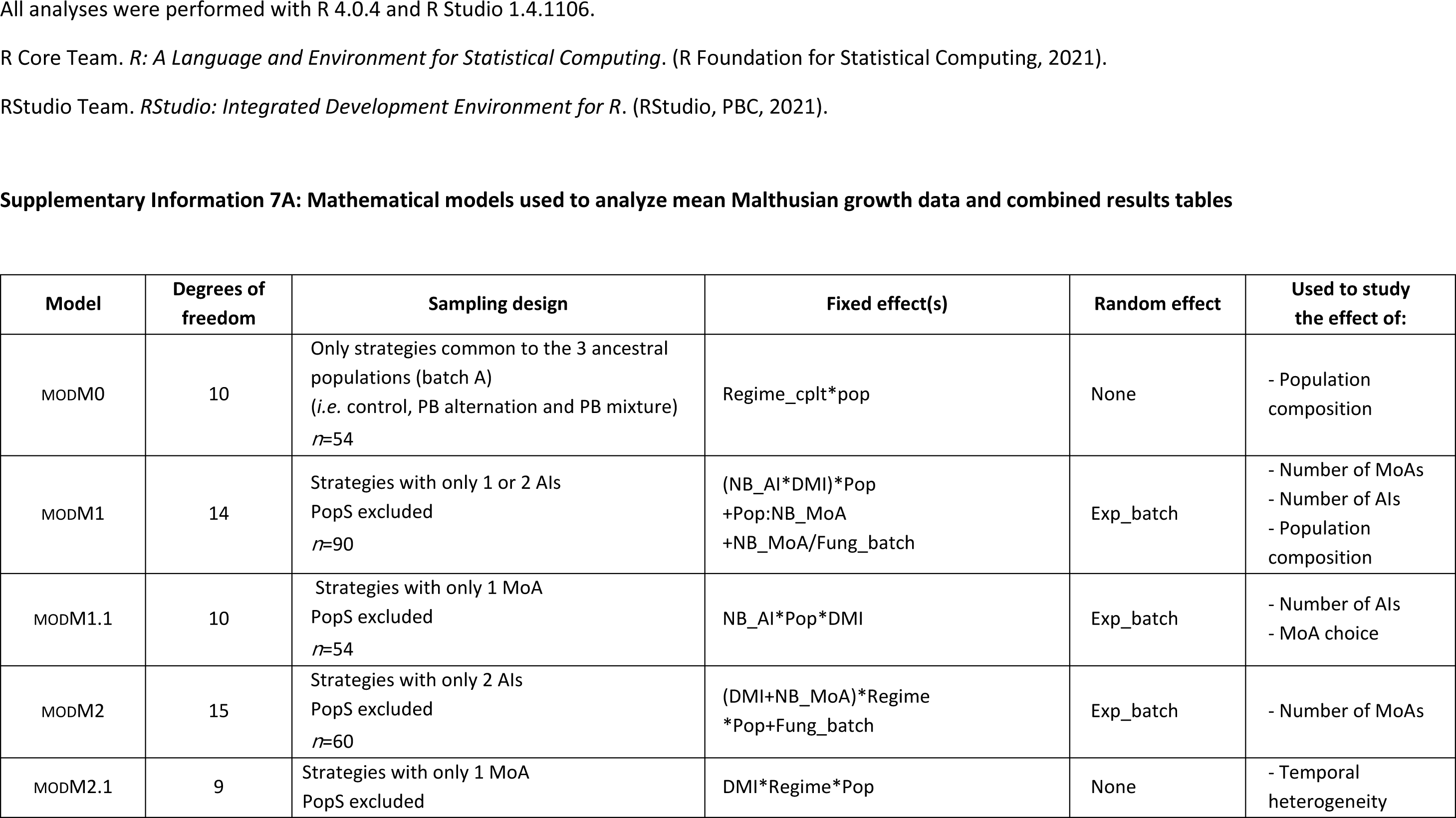
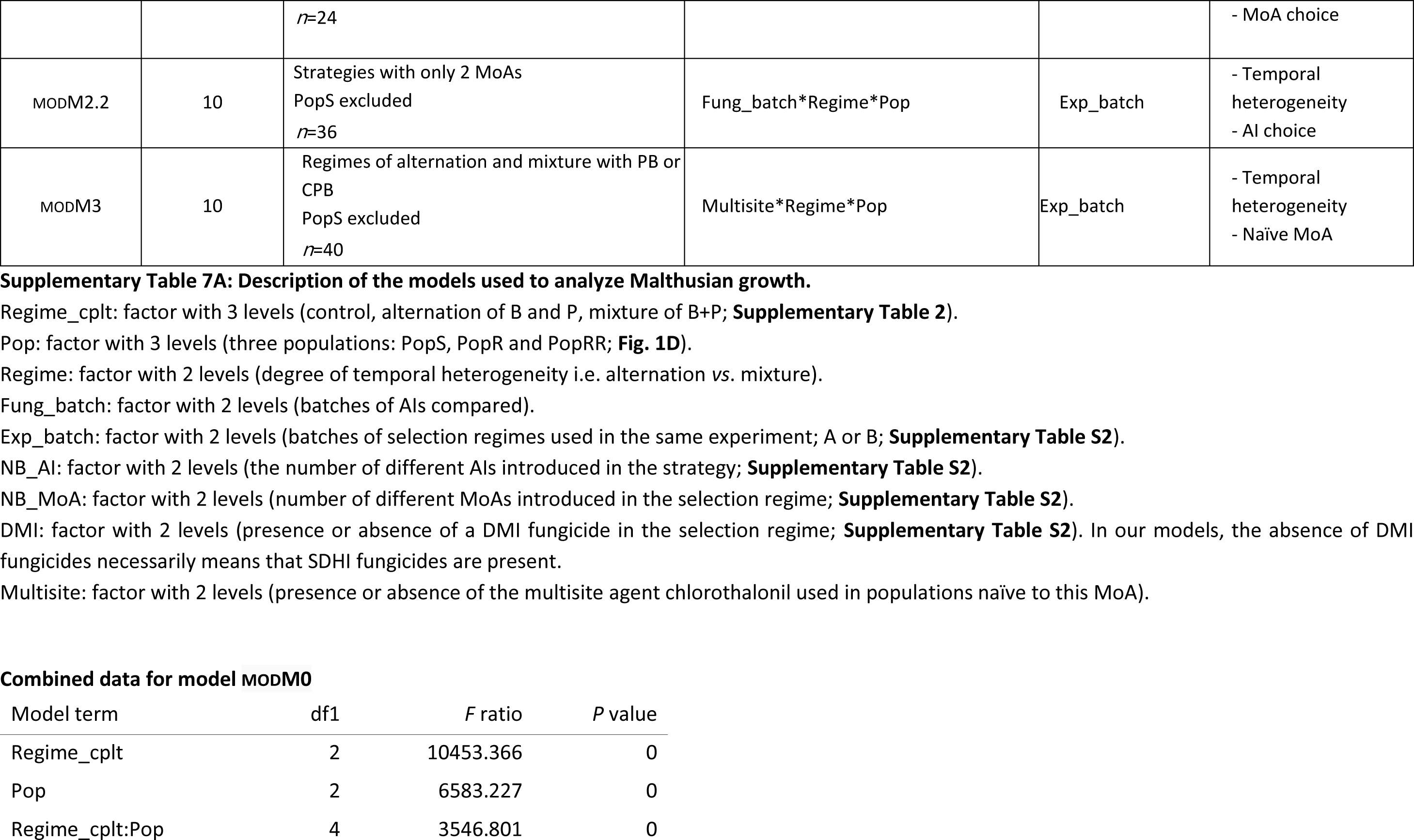
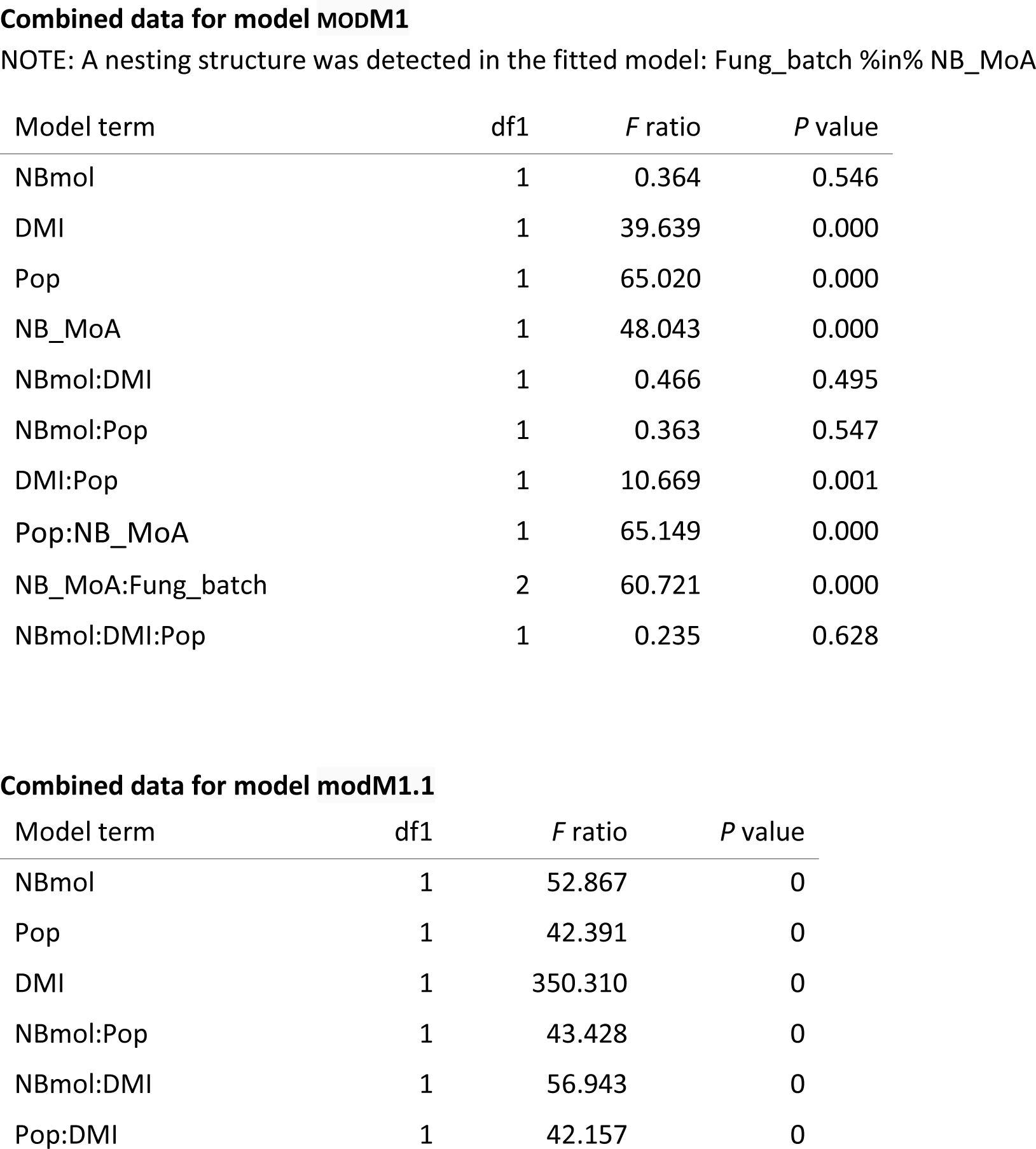
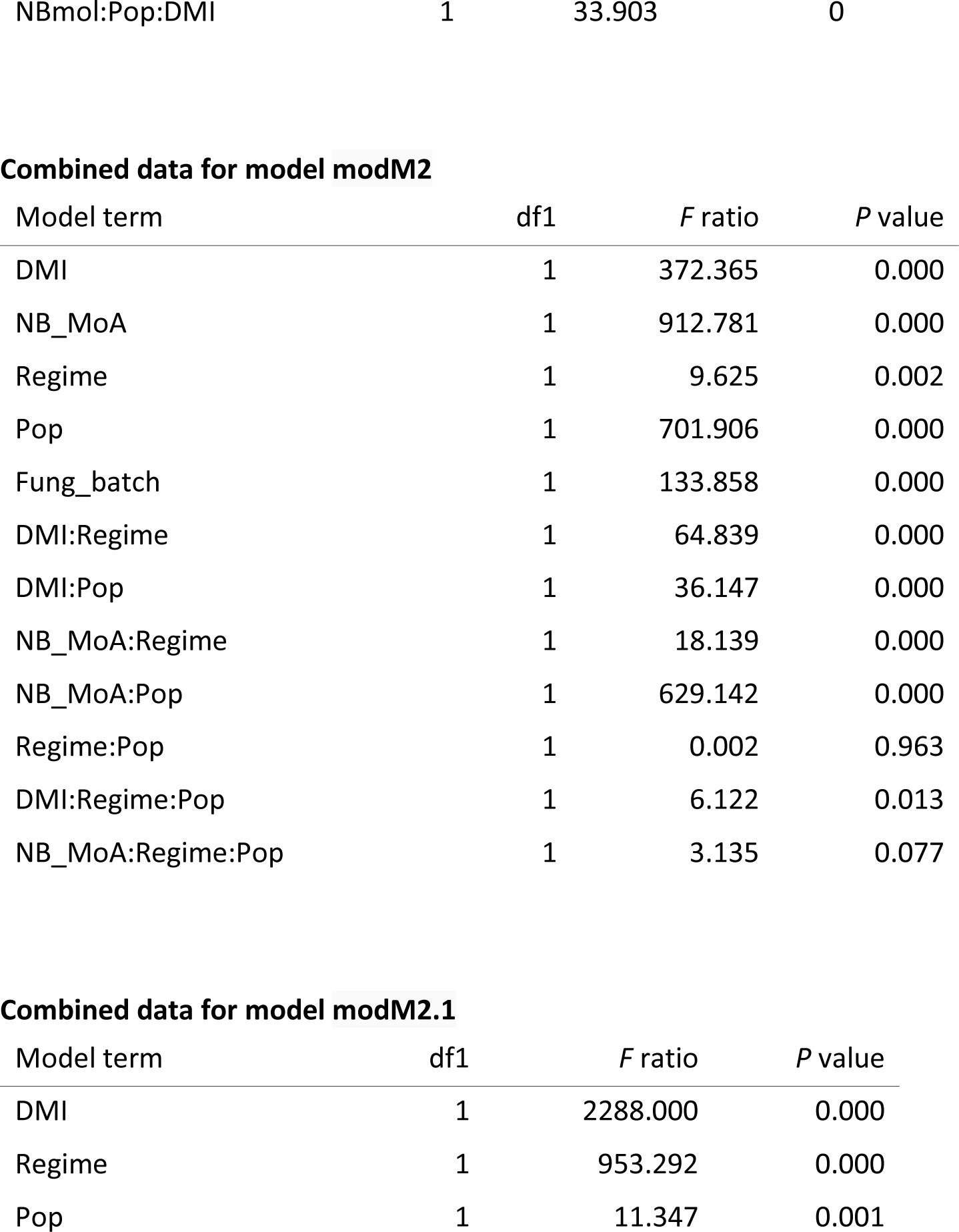
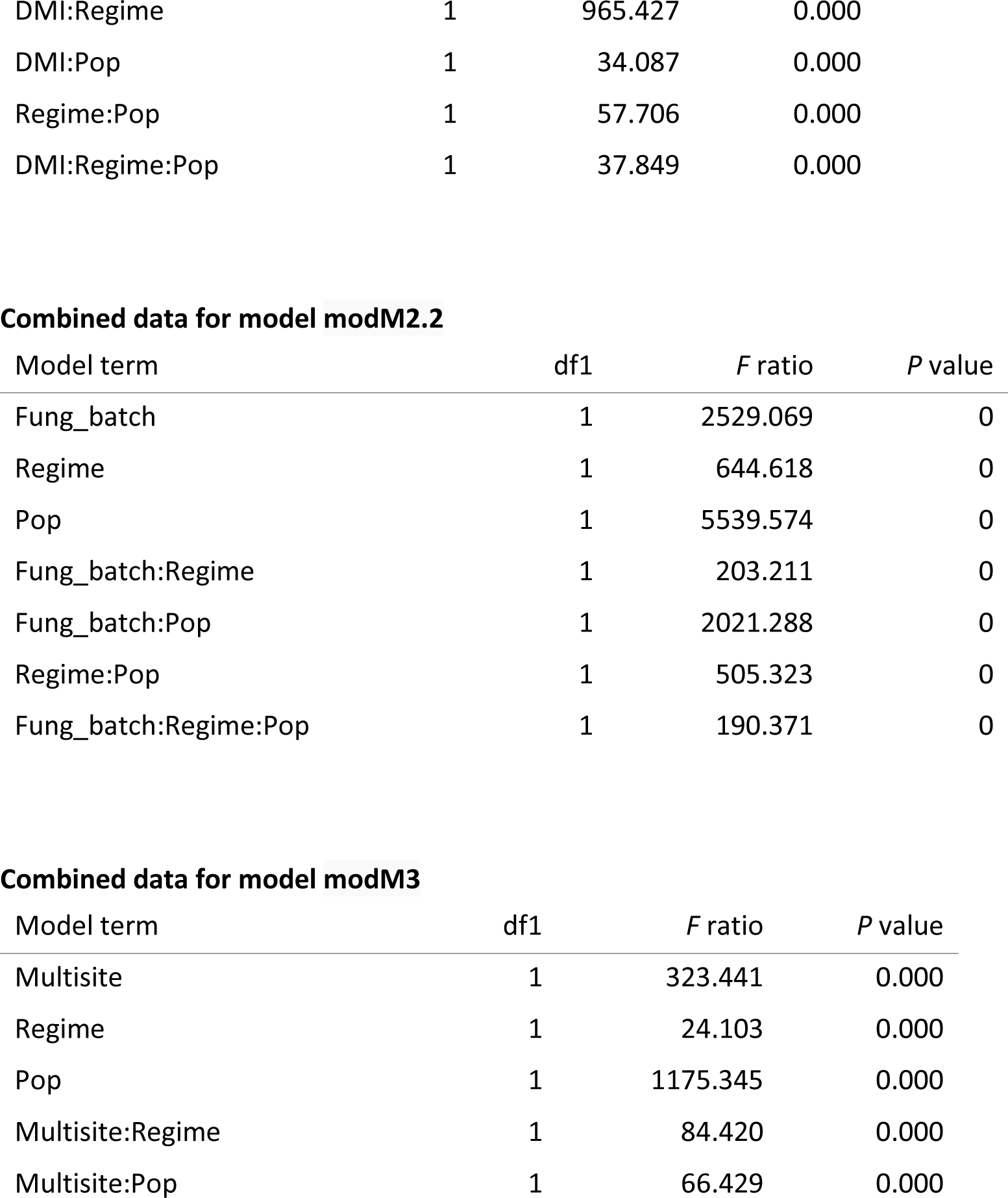
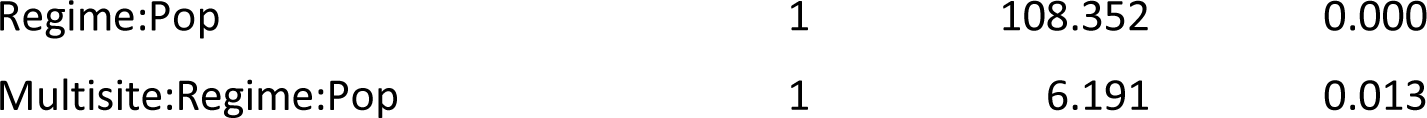
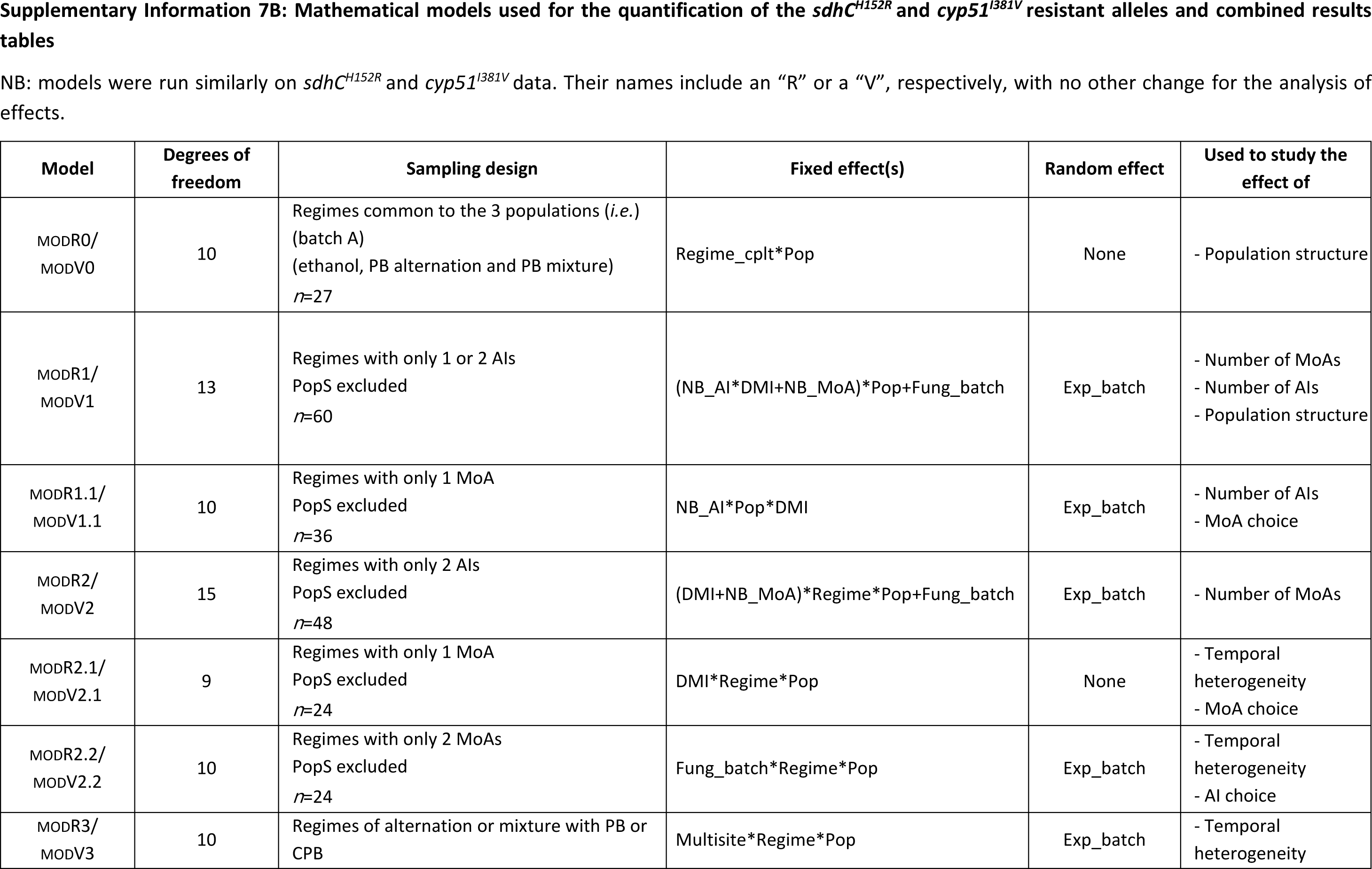
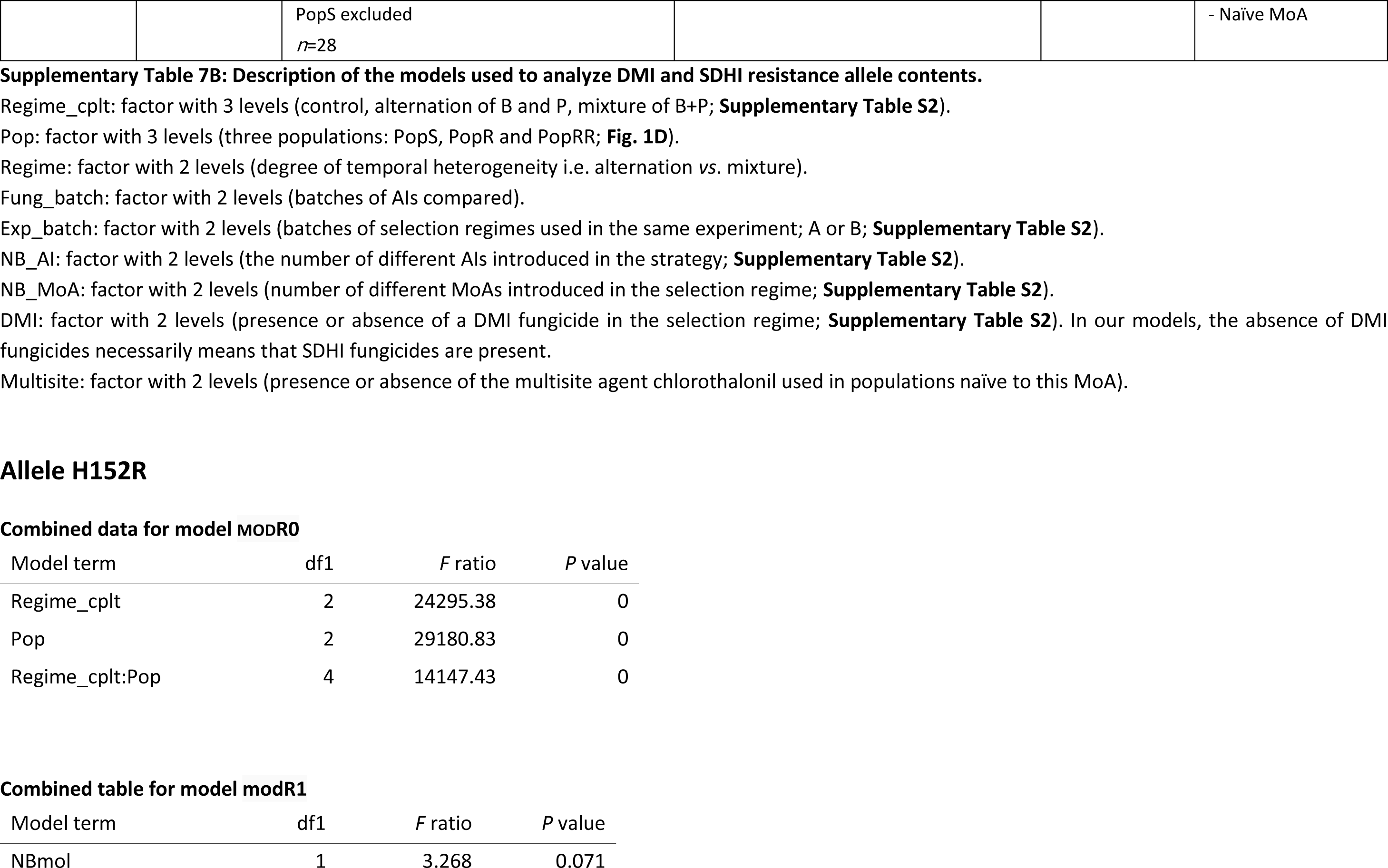
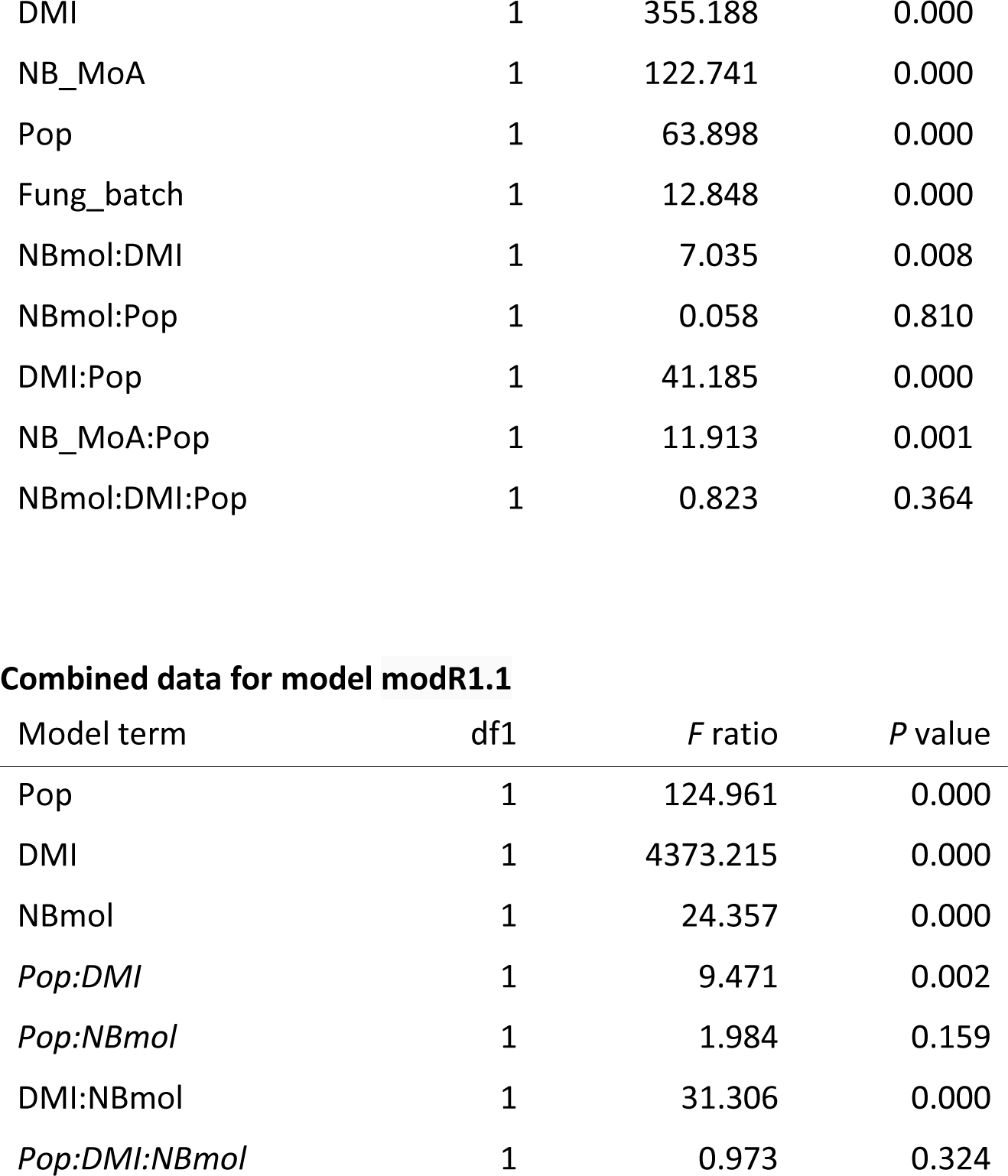
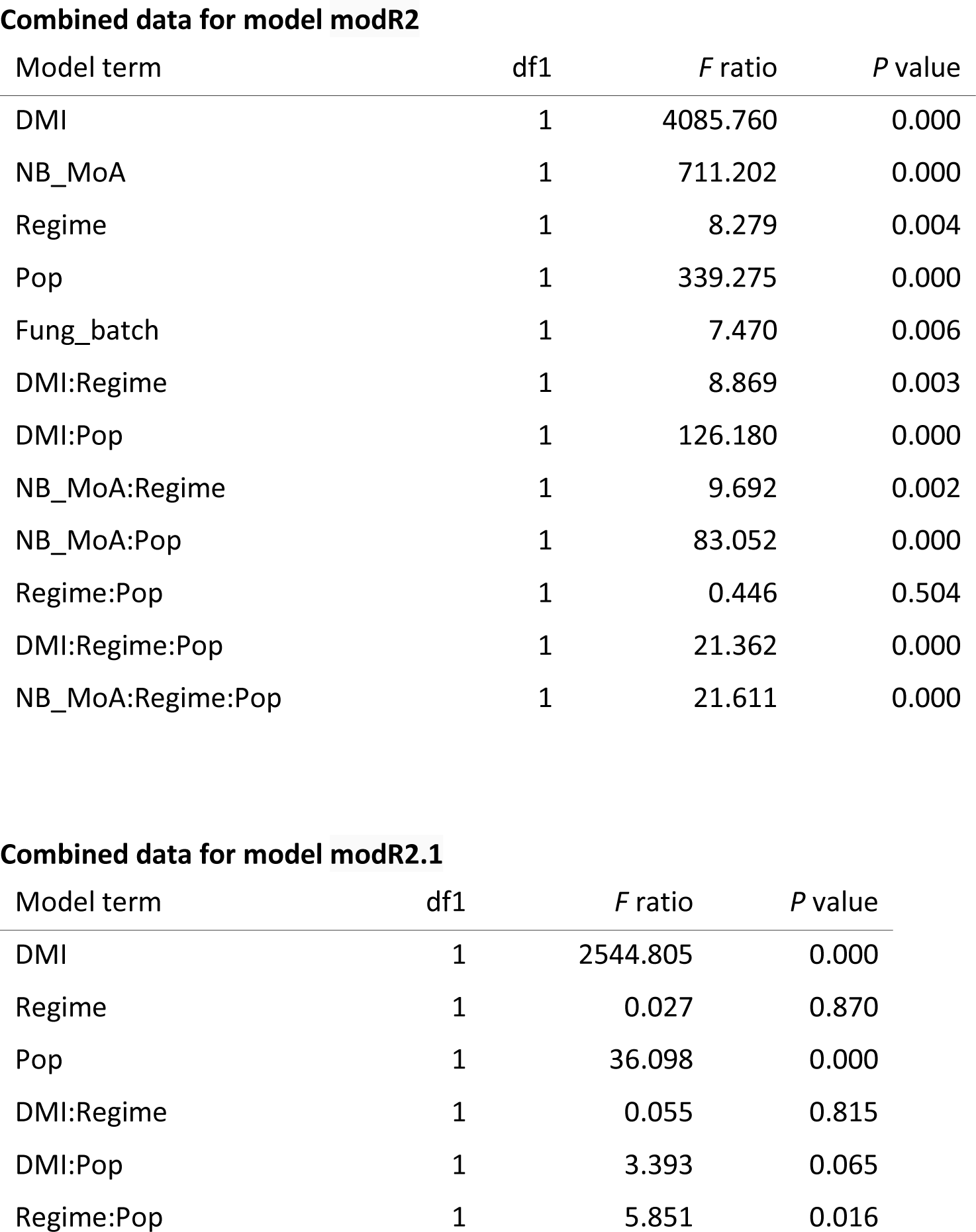
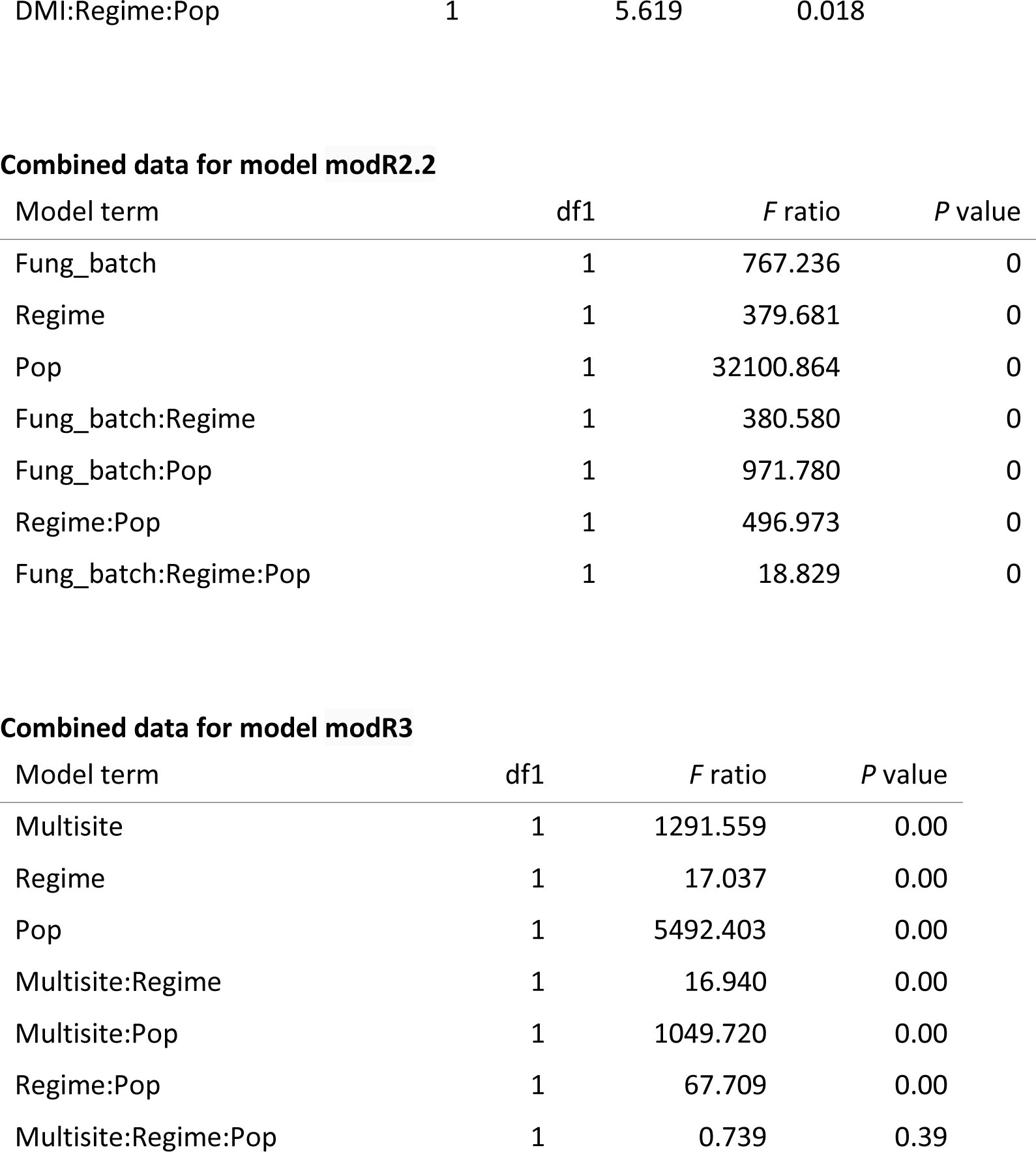
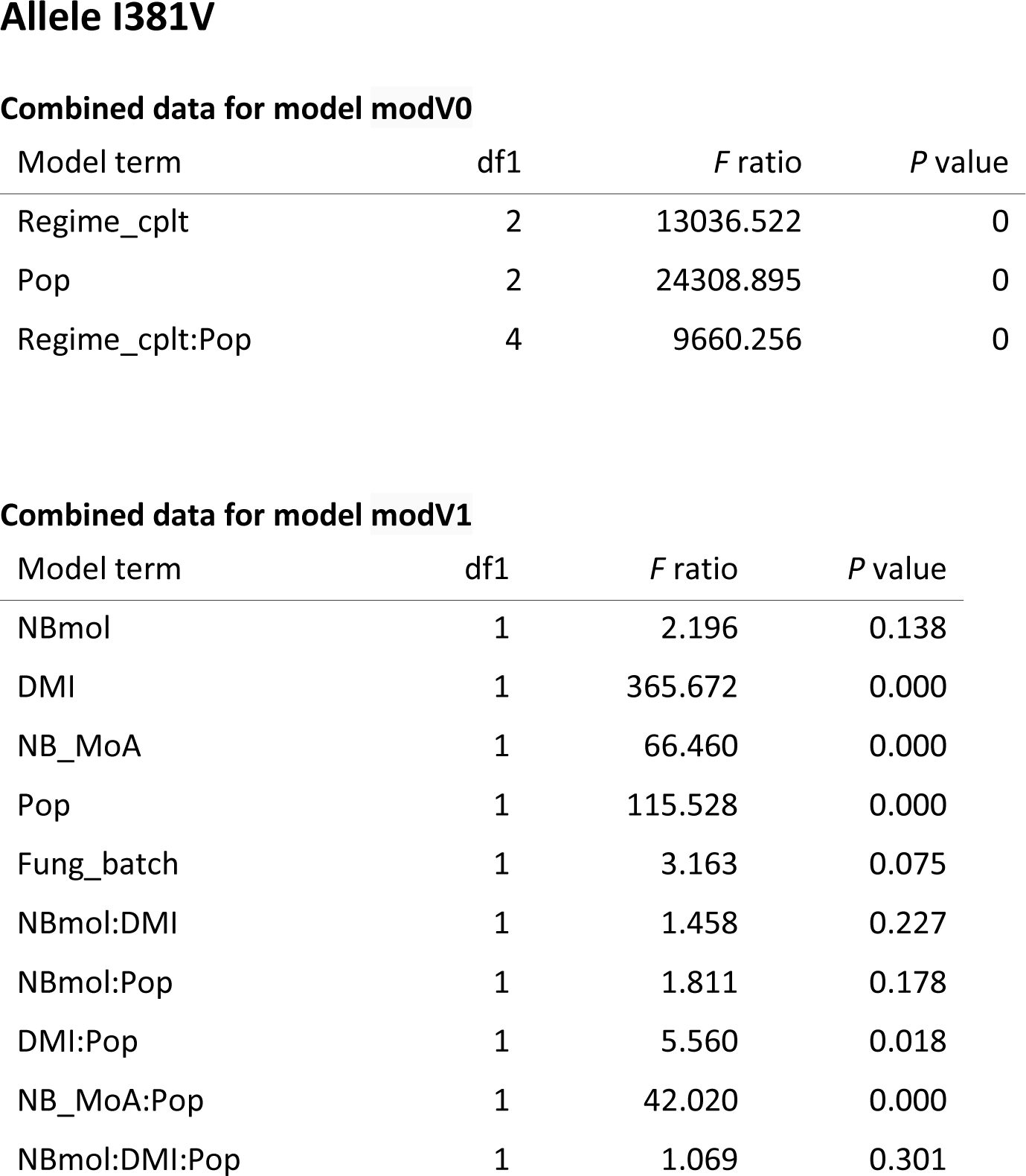
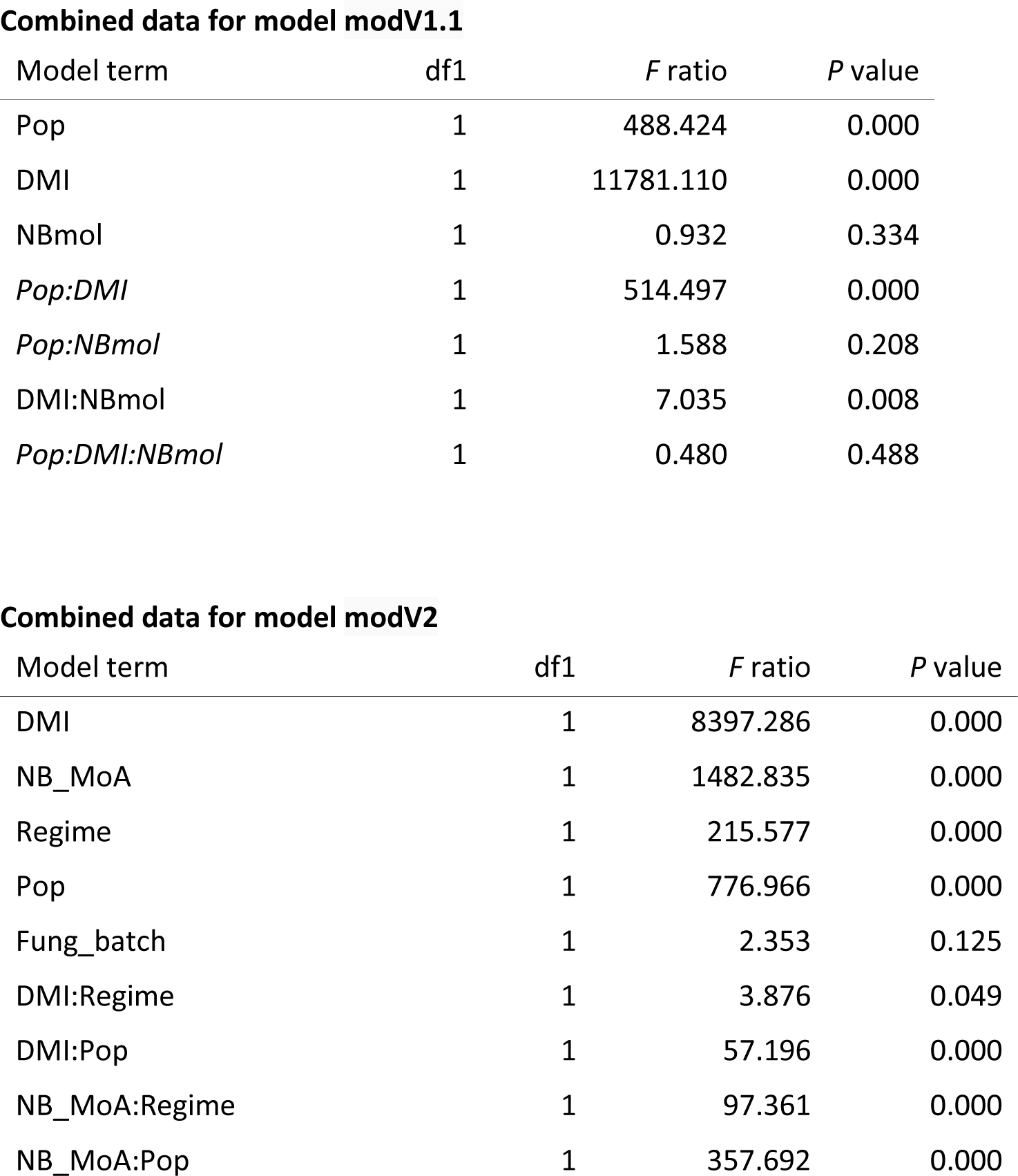
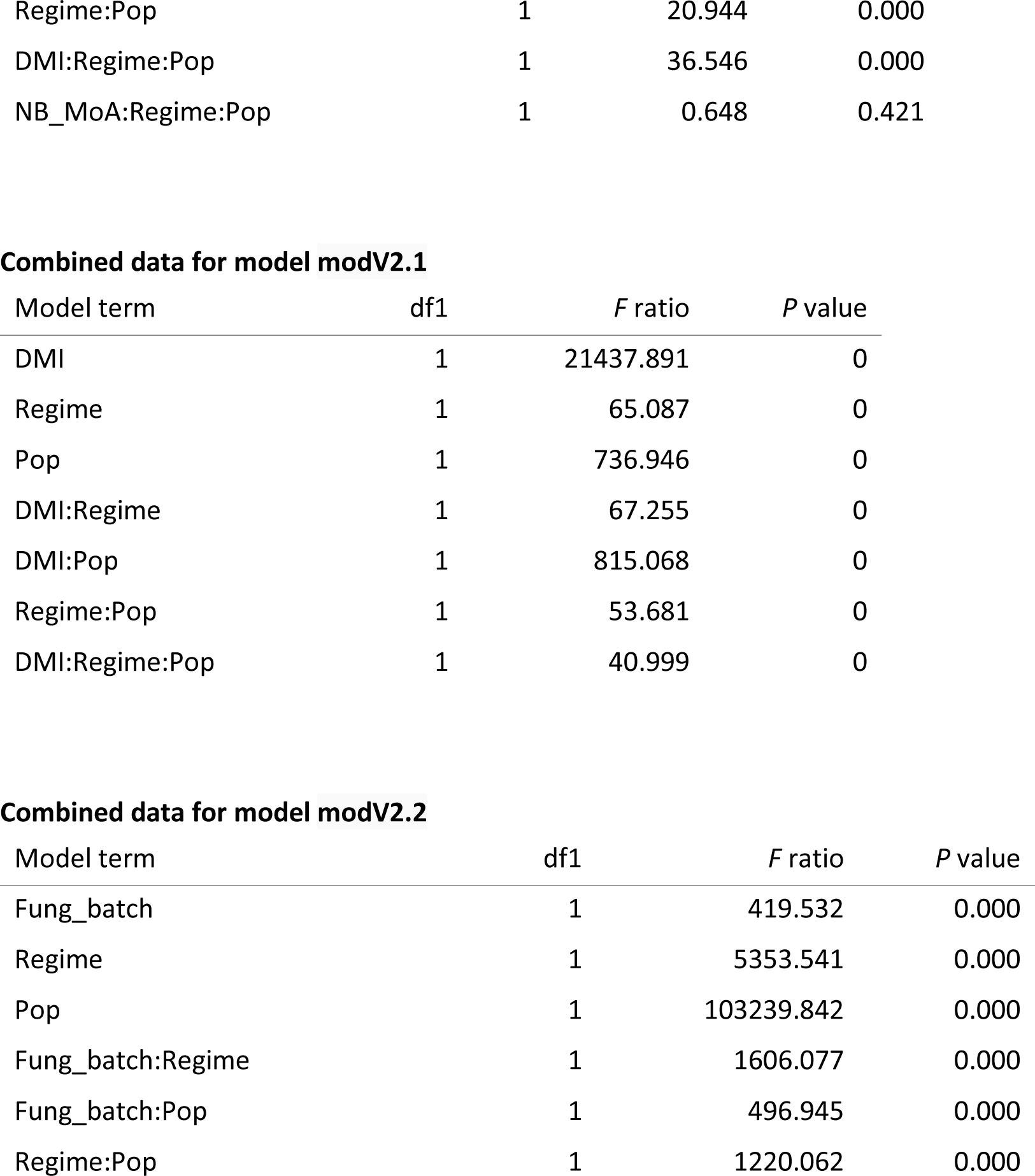
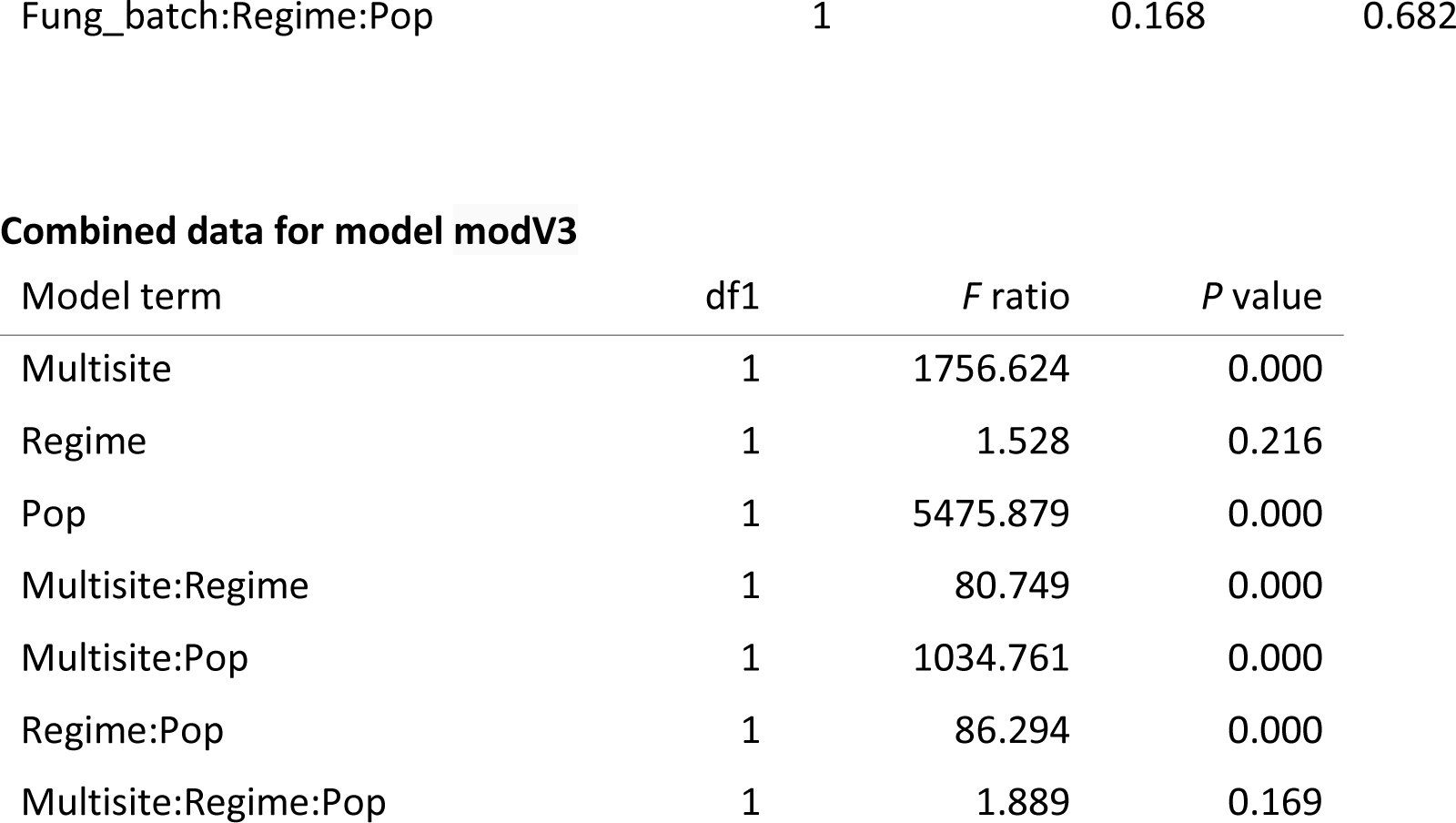
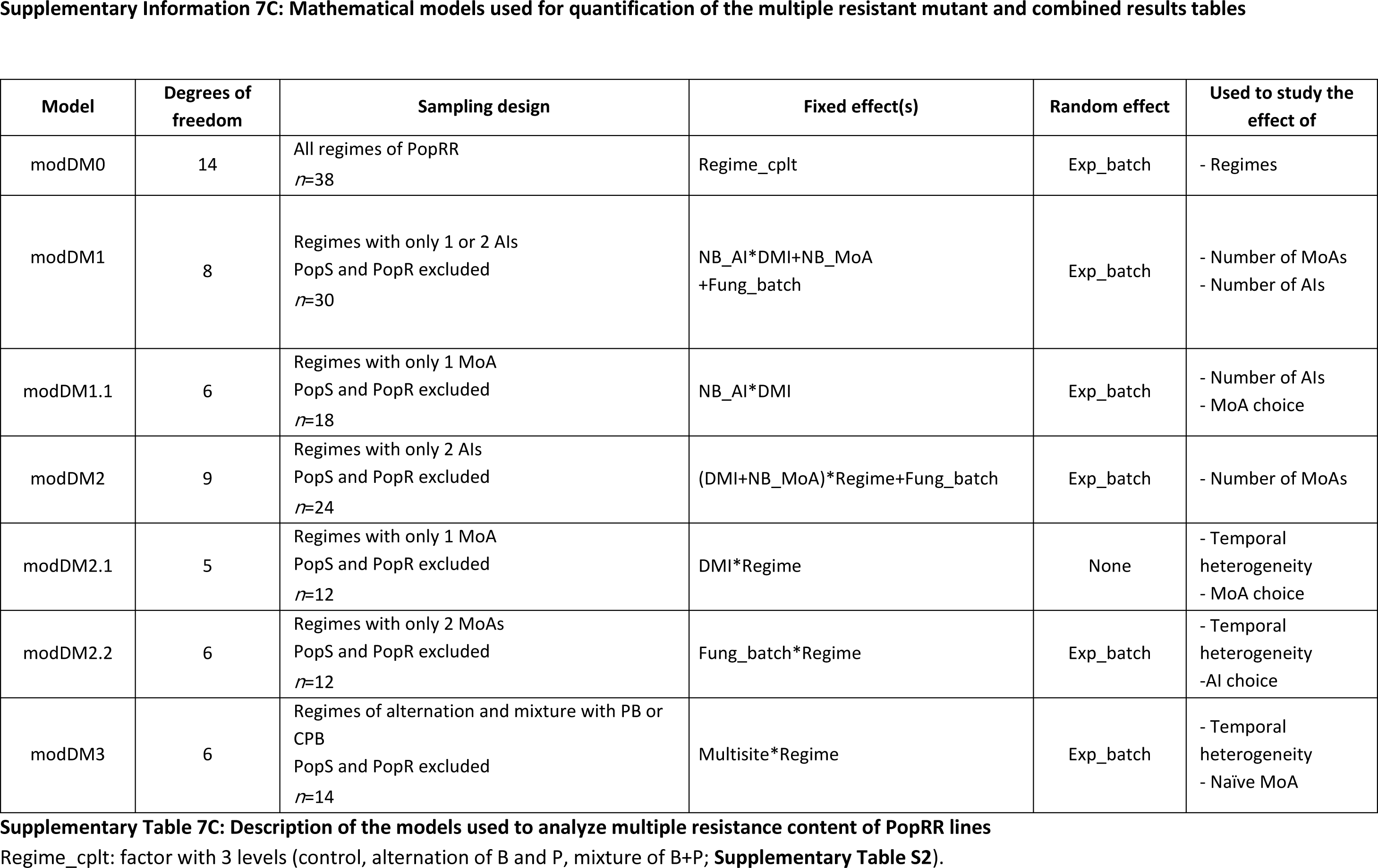
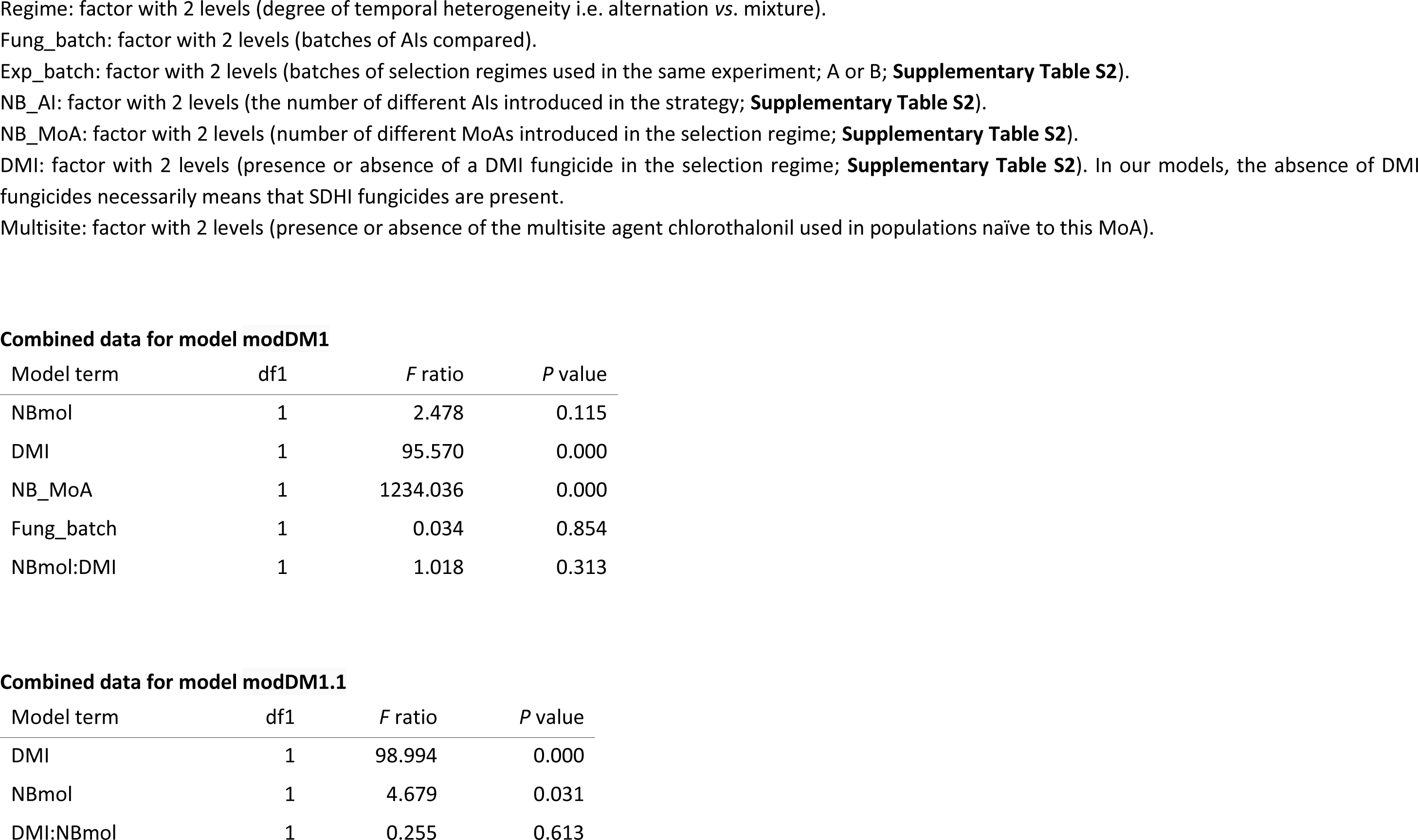
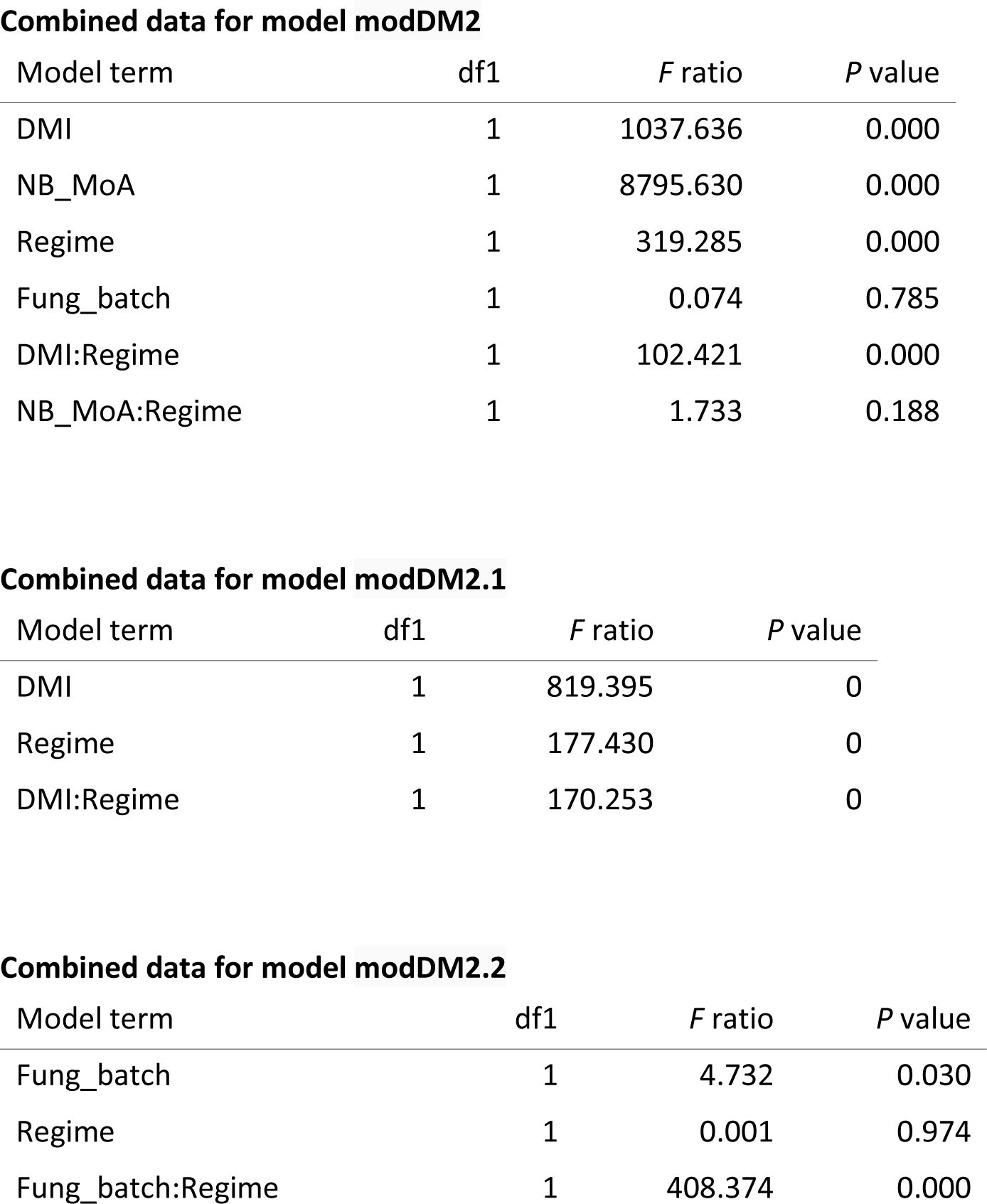
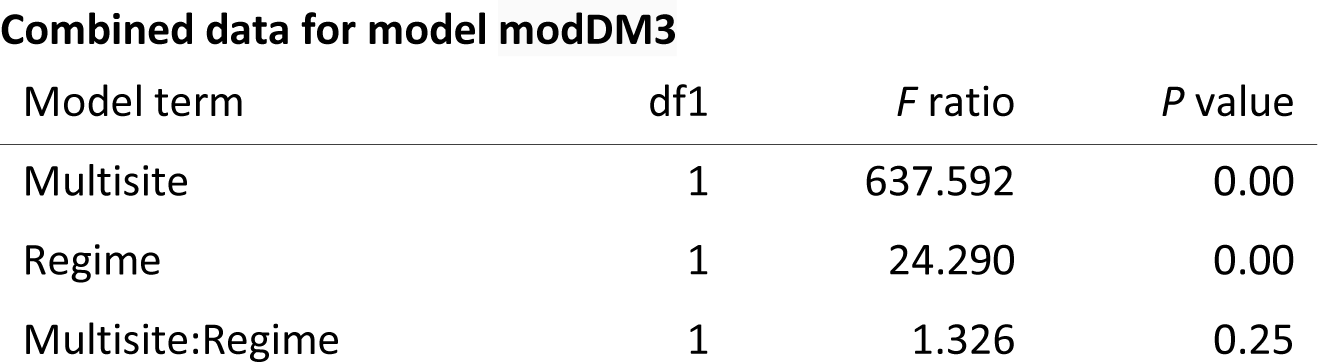
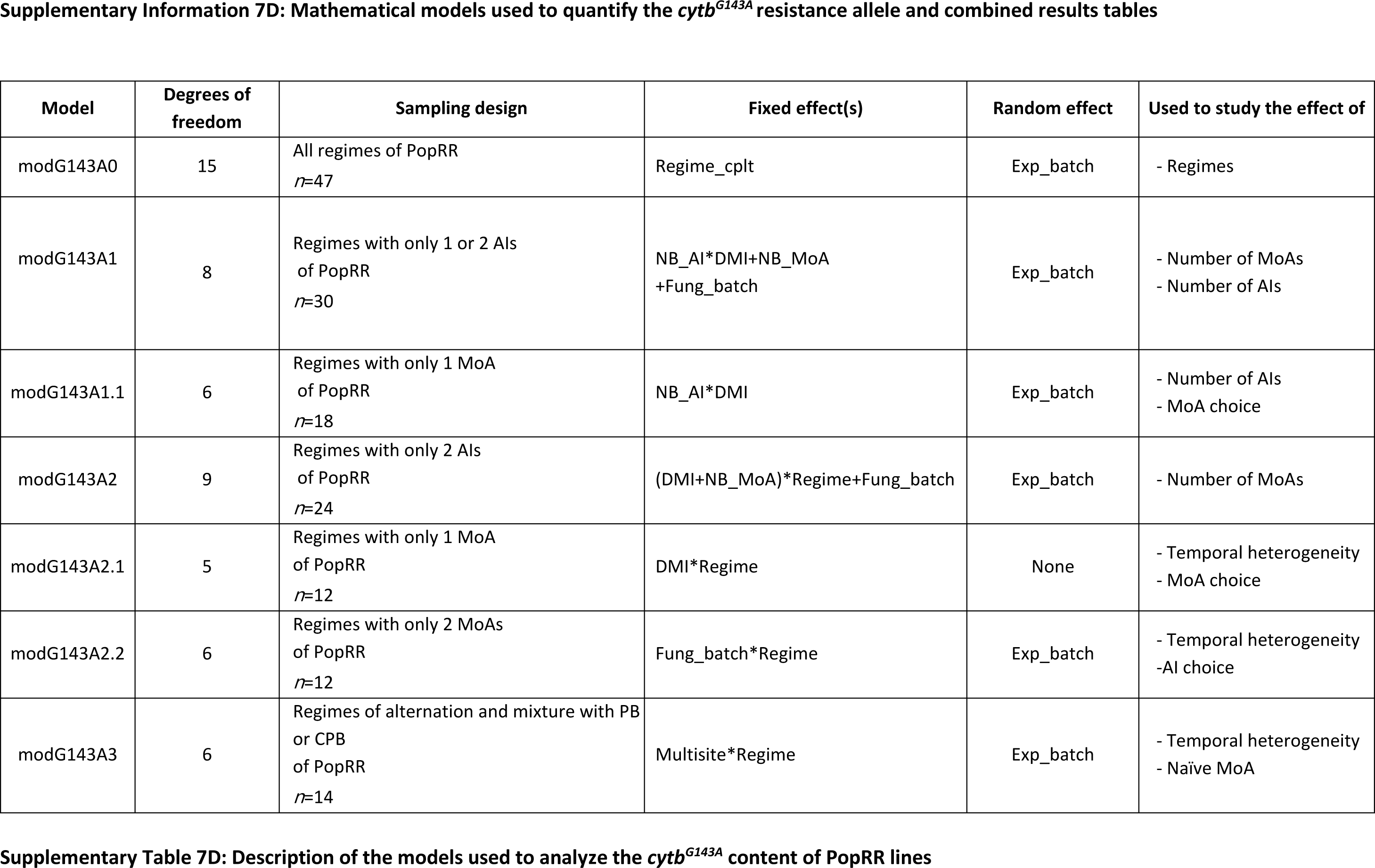
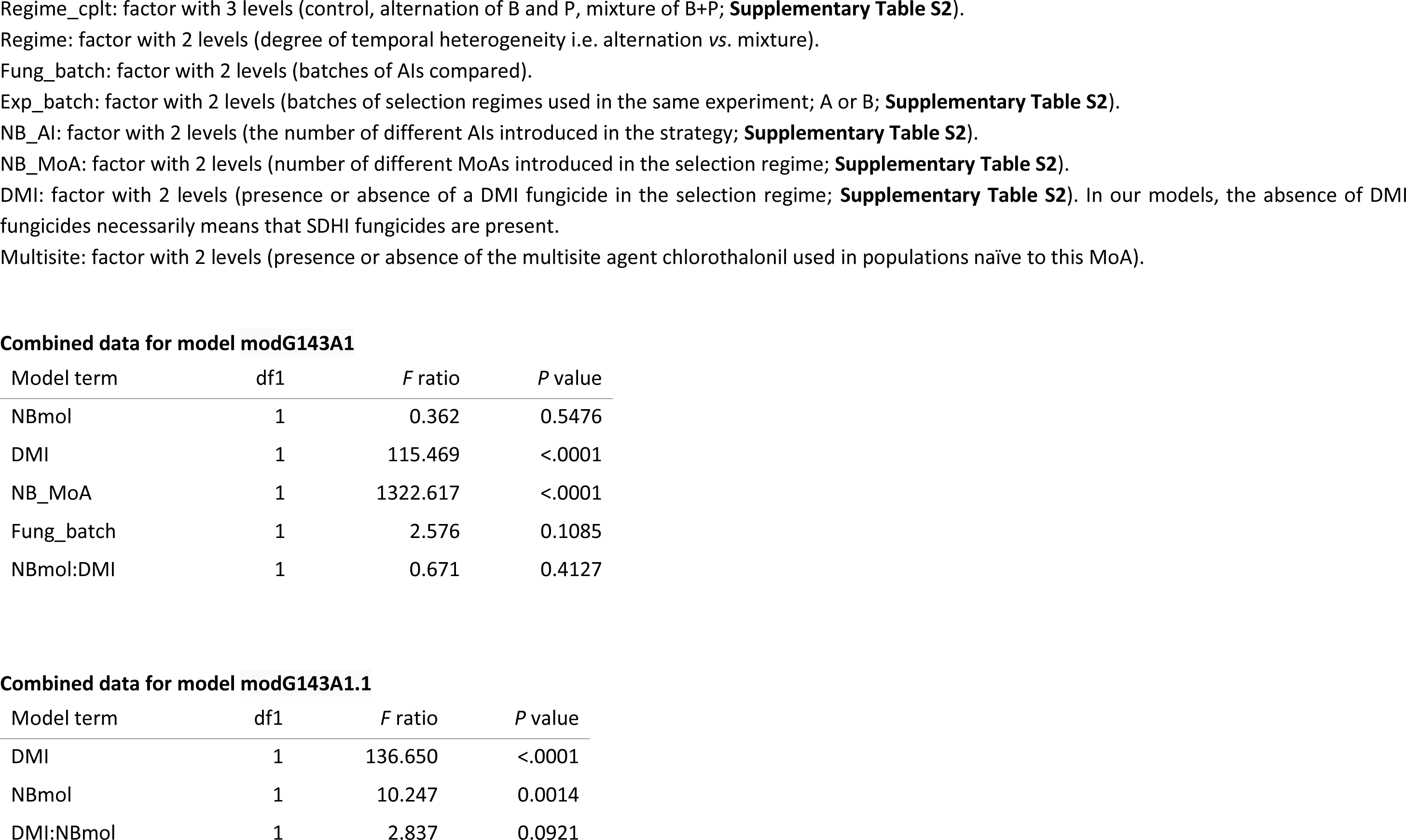
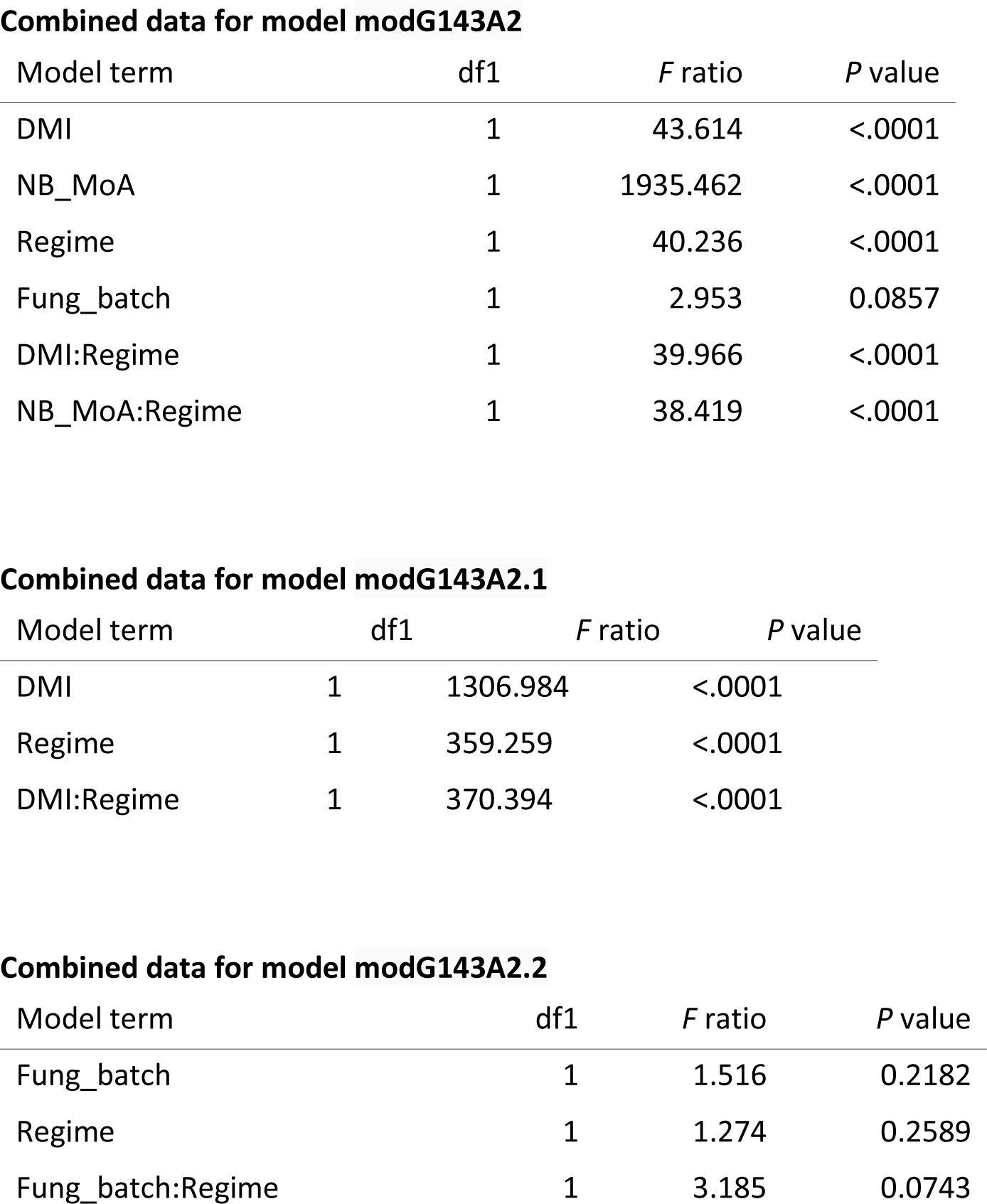
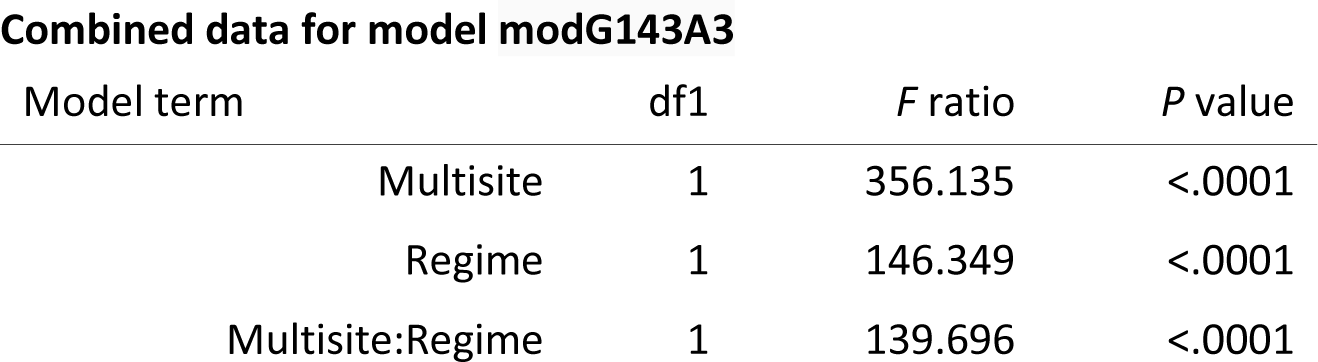

## Supplementary information 8: Performance of management strategies according to the selection regime applied to the ancestral population of *Z. tritici*

This SI is a detailed description of statistics associated to Figure 6.

***A. Performance of strategies with poorly diverse selection pressure.*** Increasing the number of AIs within a strategy did not significantly improve Malthusian growth (MODM1, *P*=0.5464), normalized resistance allele concentration (MODR1, *P*=0.0706; MODV1, *P*=0.1384) or the normalized concentration of the multiple mutant (MODDM1, *P*=0.1155). In particular, combining AIs sharing the same MoA (SDHI or DMI) did not reduce the growth of the population, actually even increasing it (MODM1.1, *P*=0.9358 for DMIs and *P*<0.0001 for SDHIs), whilst also accelerating the selection of single resistance alleles (MODR1.1, *P*<0.0001; MODV1.1, *P*=0.0236) in PopR lines, and limiting the selection of multiple resistance in PopRR lines, particularly under DMI exposure (MODDM1.1, *P*=0.3022). DMI-based strategies resulted in 1.04 times higher mean normalized Malthusian growth than SDHI- based strategies, whatever the initial population composition (PopR and PopRR, MODM1.1, *P*<0.0001), the number of AIs (1 or 2, MODM1.1, P<0.0001) or the temporal pattern of fungicide application (alternation and mixture, MODM2.1, *P*<0.0001). In PopRR lines, SDHIs controlled the multiple-resistant isolate less efficiently than DMIs (MODDM1.1, *P*<0.0001). The temporal pattern of fungicide application had little or no effect on Malthusian growth when AIs with the same MoA were combined (MODM2.1, *P*<0.0001 for SDHIs only, *P*=0.7783 for DMIs only) and on the selection of single resistance alleles (MODR2.1, in the presence of SDHIs P=0.7833, in the presence of DMIs only *P*=0.9597; MODV2.1, in the presence of DMIs only *P*=0.8505).

***B. Performance of strategies with highly diverse selection pressure.*** Combining AIs with different MoAs (SDHIs and DMIs) significantly reduced growth (modM1, *P*<0.0001) and the selection of resistance alleles (MODR1, *P*<0.0001; MODV1, *P*<0.0001) when only single-resistant isolates were introduced into the initial population. This benefit was not observed in PopRR lines (MODM1, *P*=0.2287), due to the early and rapid selection of the multiple resistant ancestral strain as opposed to its counterselection in the presence of a single MoA (MODDM1, *P*<0.0001; MODG143A1, *P*<0.0001). Strategies combining P and B were more efficient than those combining M and F for decreasing Malthusian growth (MODM2.2, *Ps*<0.0001) or the selection of single resistance alleles (MODR2.2 or MODV2.2, *Ps*<0.001) in PopR lines, and, to a much lesser extent, in PopRR lines (MODDM2.2, *P*= 0.0296). The addition of a previously unused AI (C) maintained (PopR lines; MODM3, *P*<0.0001) or restored (PopRR lines; MODM3, *P*<0.0001) the performance of strategies, but the utility of this driver depended strongly on its dose and redundancy in the strategy in which both B and P lost efficacy due to the establishment of multiple resistance (PopRR lines). Mixtures outperformed alternation when SDHIs and DMIs were combined (Malthusian growth MODM2.2, P<.001) but only if exclusively single-resistant isolates were present in the initial population. Mixture and alternation strategies performed similarly otherwise, but with alternation limiting the Malthusian growth of the multiple-resistant isolate more efficiently when a third previously unused MoA was added to SDHIs and DMIs (MOD3, *P*<0.001).

